# Spatiotemporal localization and activity of mitochondria regulate actomyosin contractility in *Drosophila* gastrulation

**DOI:** 10.1101/2025.09.05.674142

**Authors:** Somya Madan, Sayali Chowdhary, Sakshi Phalke, Harsh Mittal, Manos Mavrakis, Richa Rikhy

## Abstract

Mitochondrial dynamics are vital for embryogenesis, but their role in morphogenesis is largely unexplored. In this study, we elucidate the role of mitochondrial dynamics and function in regulating apical constriction at the ventral furrow in *Drosophila* gastrulation. We find that spatiotemporal redistribution of mitochondria from basal to apically constricting regions is coincident with Myosin II accumulation in ventral epithelial cells. This apical enrichment depends on intact microtubules and is regulated by Dorsal/NFkb signaling pathway. Embryos expressing mitochondrial fission protein, Drp1 mutants, contain larger, basally accumulated mitochondria, decreased apical constriction and ventral furrow defects. Transcriptomic and proteomic profiling of Drp1 mutants shows no change in Dorsal signaling, but an upregulation of antioxidant enzymes, decreased respiratory complexes, and reduced reactive oxygen species (ROS), confirming loss of mitochondrial activity and a reduced environment. Depleting mitochondrial SOD2 in Drp1 mutant embryos restores levels of antioxidant proteins and increases ROS, leading to an oxidized environment. These embryos also show rescue in apical mitochondrial migration and Myosin II accumulation, thereby enabling proper ventral furrow formation. Our study reveals the functional importance of localized mitochondrial dynamics, regulated by the Dorsal signaling pathway, and mitochondrial activity in a spatiotemporal manner for apical constriction during gastrulation.

## Introduction

Embryonic development is elaborate and dynamic, involving various morphogenetic processes and cellular remodeling events such as cell division, migration, tissue folding and invagination. These intricate cellular movements occur due to feedback between signaling pathways and organelle function. They are energy-intensive and require specific metabolites for appropriate activation. Mitochondria, the powerhouse of the cell, are the source of energy and metabolites essential for supporting the dynamic morphogenetic events during embryogenesis (Dumollard et al., 2009; Madan et al., 2021; May-Panloup et al., 2021). Apart from ATP production, mitochondria have multiple other functions, such as redox balance, calcium buffering and epigenetic regulation through which they can modulate signaling pathways and cytoskeletal architecture. Mitochondria are well known to be integral to morphogenetic processes such as epithelial to mesenchymal transition (Zhang et al., 2020), cell division (Finkel and Hwang, 2009; Martínez-Diez et al., 2006; Mitra et al., 2009), cell migration (Kim et al., 2015; Lim et al., 2015; Wang et al., 2015) and wound healing (Fu et al., 2020; Xu and Chisholm, 2014). Mitochondrial distribution to sites of active cellular dynamics and local release of metabolites aids in driving cell shape changes.

Mitochondria are maternally inherited in most organisms; hence, oocyte mitochondrial health is a crucial determinant of proper embryonic growth and development (Giles et al., 1980). Oocyte and early embryo mitochondria are small and spherical with low ATP production and oxygen consumption (Chowdhary et al., 2020, 2017; Cox and Spradling, 2003; Dumollard et al., 2009; Jonathan and Davis, 1995). In humans, it has been shown that oocytes with mitochondria exhibiting low electron transport chain (ETC) activity can lead to defective embryonic development (Heng-Kien et al., 2005; Wilding et al., 2001). Disruption of mitochondrial membrane potential and ETC complexes causes developmental arrest in zebrafish (Arribat et al., 2019).

Mitochondria are dynamic and undergo regulated cycles of fusion and fission, resulting in varied morphologies that range from globular to intricate reticular networks. The Dynamin superfamily of large GTPases carries out mitochondrial fusion and fragmentation. Dynamin-related protein 1 (Drp1) functions in mitochondrial fission, by assembling around the mitochondrion, constricting it and separating it into two daughter mitochondria via its GTPase activity (Bleazard et al., 1999; Labrousse et al., 1999; Sesaki and Jensen, 1999). Mitochondrial fusion is carried out by Mitofusins 1 and 2 (Mfn1 and 2) and Optic atrophy 1 (Opa1). Mfn1 and 2 fuse the outer mitochondrial membrane, and Opa1 is responsible for inner mitochondrial membrane fusion (Ban et al., 2018; Hsiuchen et al., 2003; Olichon et al., 2003). Mitochondrial fission enables inheritance of functional mitochondria into the oocyte(Lieber et al., 2019). Mitochondrial fission and fusion play a crucial role in embryonic development. Mfn1/2 mutant mouse embryos are embryonic lethal (Hsiuchen et al., 2003). Loss of Opa1 in mouse embryos also leads to retarded growth and embryonic lethality (Moore et al., 2010). Maternal depletion of Drp1 and Opa1 in *Drosophila* leads to a decrease in embryo formation and lethality (Chowdhary et al., 2020; Mitra et al., 2012). Drp1 knockout mice are also embryonic lethal, and they exhibit neurological defects (Ishihara et al., 2009; Wakabayashi et al., 2009). Knockdown of Drp1 in *C. elegans* also causes embryonic lethality (Labrousse et al., 1999). Mitochondria also contribute to embryonic development by producing metabolites that act as regulators of gene expression (Gut and Verdin, 2013; Lees et al., 2017). Mouse embryos cultured in the presence of mitochondrial metabolic inhibitor, amino-oxyacetate, show reduced fetal and placental growth (Wakefield et al., 2011). Differential mitochondrial positioning and metabolic activity, along the embryonic axis, regulate cell fate determination in *Xenopus* and sea urchin embryos (Chang et al., 2017; Coffman, 2009; Czihak, 1963; Garfinkel et al., 2023; Yost et al., 1995). However, the mechanism by which mitochondrial morphology, activity and distribution modulate cell shape changes and morphogenesis during embryonic development remains unexplored.

*Drosophila* gastrulation is a well defined model system to study mitochondrial distribution and function in morphogenesis. It depends upon rapid cellular movements taking place at different domains of the embryo, including internalization of the mesoderm precursor cells by invagination called the ventral furrow (VF) formation. It begins with a change in the shape of a few cells at the ventral midline encompassing 10 to 12 cells in width and 80 cells in length. These cells first undergo flattening of the apical membrane, followed by apical constriction with the help of the acto-myosin contractile network, changing from columnar to wedge shape. The neighbouring cells stretch towards the ventral midline simultaneously. As a result of these two events, the constricting cells invaginate, forming a U-shaped VF which is later internalised, forming the mesoderm (Gheisari et al., 2020; Leptin and Grunewald, 1990; Martin, 2020; Turner and Mahowald, 1977). VF formation is activated by the transcription factors Twist and Snail downstream of the Dorsal/NFkB signaling pathway (Ko and Martin, 2020; Leptin and Grunewald, 1990).

We examine mitochondrial dynamics during VF formation in gastrulation (Figure 1A) in *Drosophila* embryos to understand the role of mitochondria in driving morphogenesis. Mitochondria are fragmented and highly abundant in early *Drosophila* blastoderm embryos (Chowdhary et al., 2020, 2017). We find that mitochondrial migration from basal to apical regions occurs during gastrulation in the VF cells, regulated by microtubules and the Dorsal/NFkB signaling pathway. Mitochondrial fission protein Drp1 mutant embryos show accumulation of larger mitochondria basally that fail to migrate apically, and embryos contain defects in VF formation. Mitochondria in Drp1 mutant embryos are functionally compromised with decreased ETC components and ROS levels. Drp1 mutant embryos do not exhibit changes in the Dorsal/NFkB pathway genes, but instead lead to increased expression of anti-oxidant genes. Increasing ROS genetically by depletion of mitochondrial SOD2 or decreasing mitochondrial fusion protein Opa1 in Drp1 mutant embryos results in mitochondrial fragmentation and apical migration in the VF, also rescuing the furrow defects. Thus, we propose that fragmented mitochondria migrate apically in the ventral cells during gastrulation to activate Myosin II by increasing ROS locally, thereby facilitating apical constriction and enabling proper VF formation.

**Figure 1:**
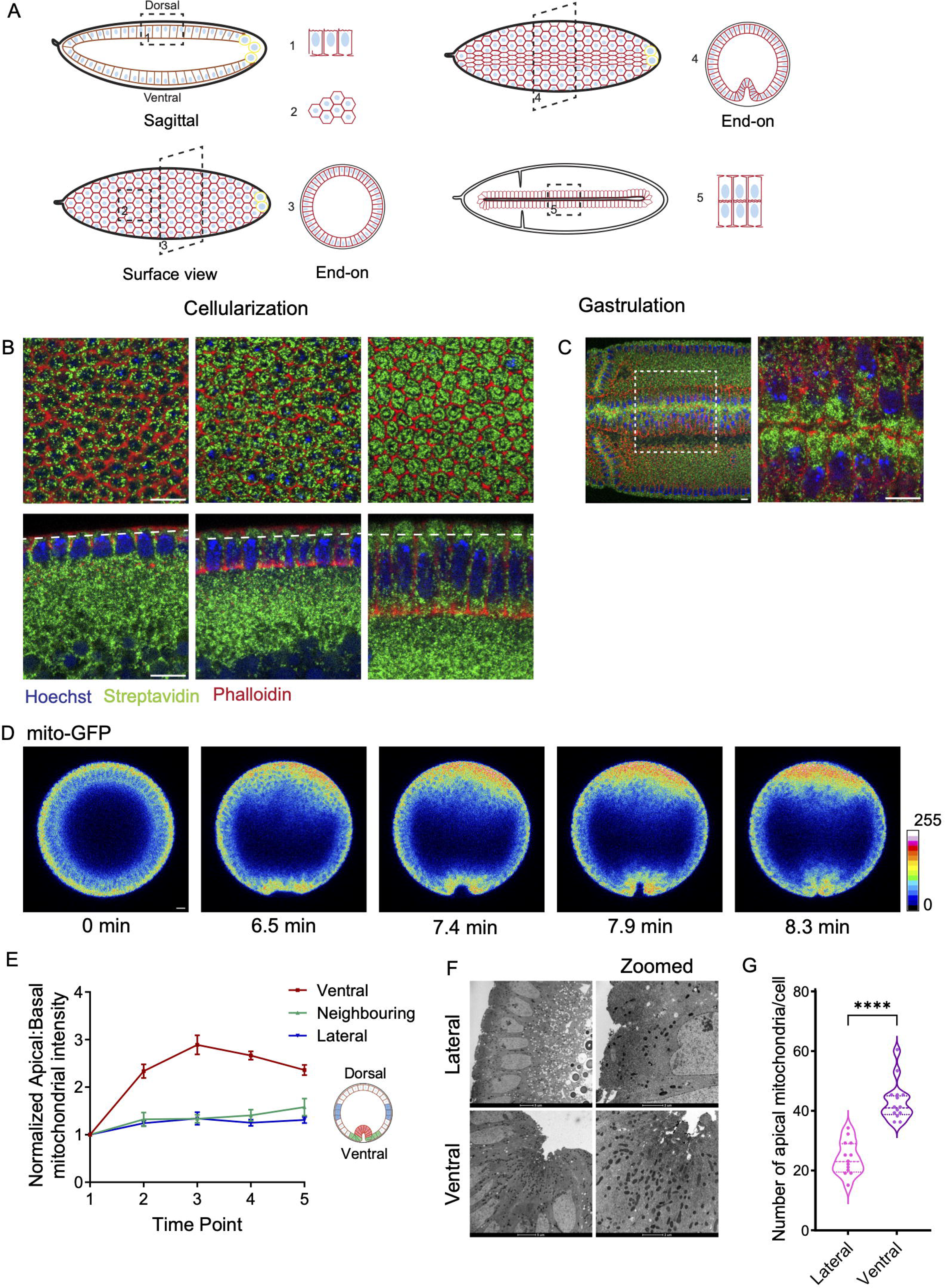
Mitochondria are enriched apically in the ventral furrow cells during gastrulation. (A) Schematic representing the *Drosophila* embryo in cellularization (left top sagittal view, left bottom surface view) and gastrulation (right) and imaging orientations (shown with box) used in fixed and live embryos to assess mitochondrial distribution and dynamics in cellularization and gastrulation. (B-C) Mitochondria are present as small punctae and enrich apically during cellularization and gastrulation. Embryos in early, mid and late cellularization (B) and in gastrulation at the VF (C) are labelled for mitochondria (fluorescent streptavidin, green), F-actin (fluorescent phalloidin, red) and DNA (Hoechst, blue). (D) Representative snapshots from live embryos expressing mito-GFP in late cellularization (first time point) and during VF formation in gastrulation. Dorsal is up and ventral is down. The images are shown in 16 color rainbow scale on an 8 bit image where 0 is black and white is 255. (E) The plot shows apical:basal mean fluorescence intensity of mito-GFP quantified in three regions of the gastrulating embryos, ventral cells (red), neighbouring cells (green) and lateral cells (blue), across 5 time points (n=3). The schematic of gastrulating embryo shows the regions chosen for quantification. Data is represented as Mean ± SEM. (F) Representative TEM images from gastrulating embryos are shown in the lateral and ventral region. Mitochondria are dark punctate structures enriched in apical regions. Zoomed in images show mitochondria enriched in apical regions of ventral cells compared to lateral cells. (G) The plot shows quantification of the number of mitochondria above the nucleus per cell in ventral and lateral cells (n= 14 and 13 cells, respectively from 4 embryos). The violin plot shows median and quartiles. The statistics were performed by two-tailed unpaired Mann-Whitney test, ****p<0.0001 Scale Bar: 10μm (B, C, D), 5μm (F).

## Materials and methods

### Key resources table

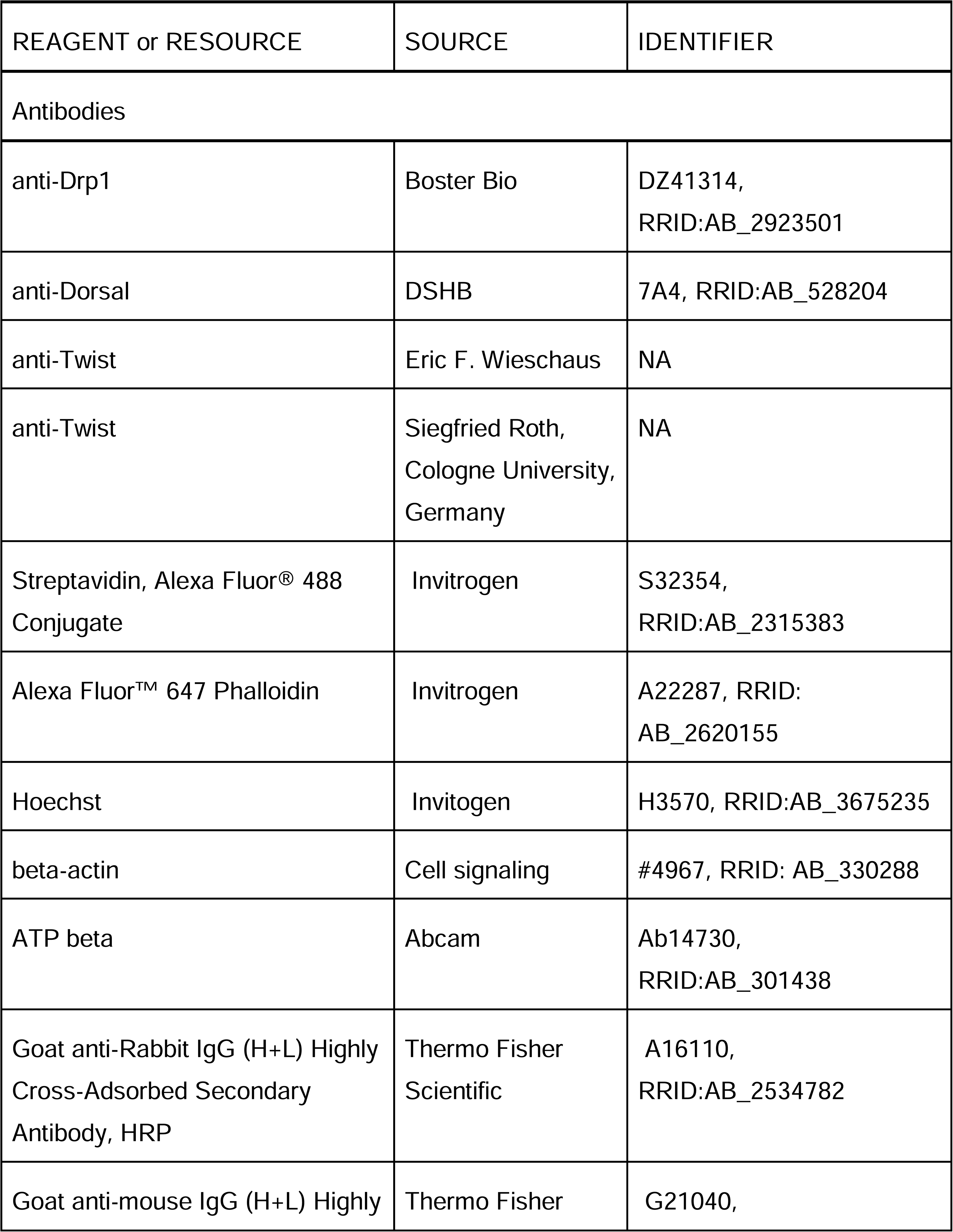
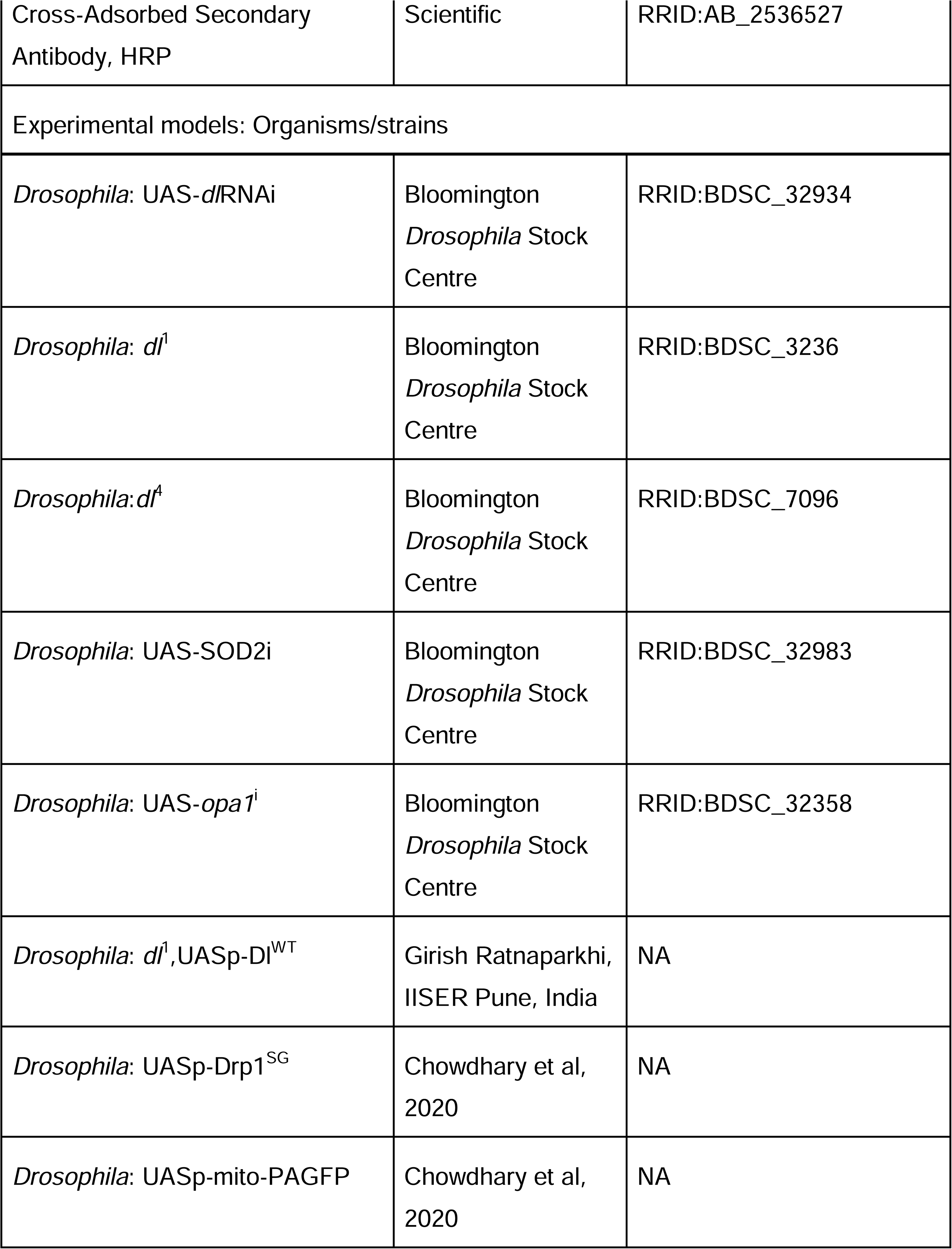
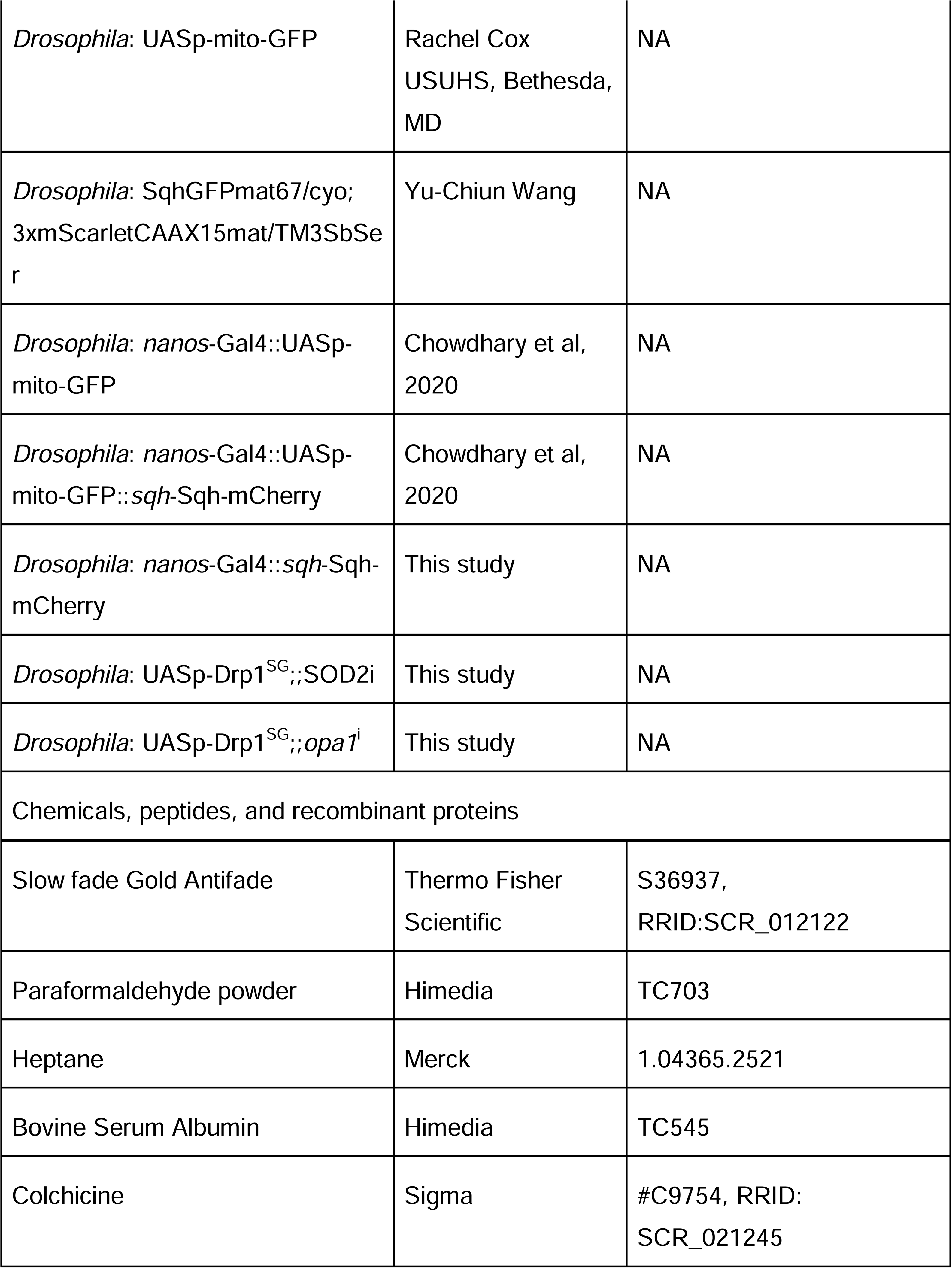
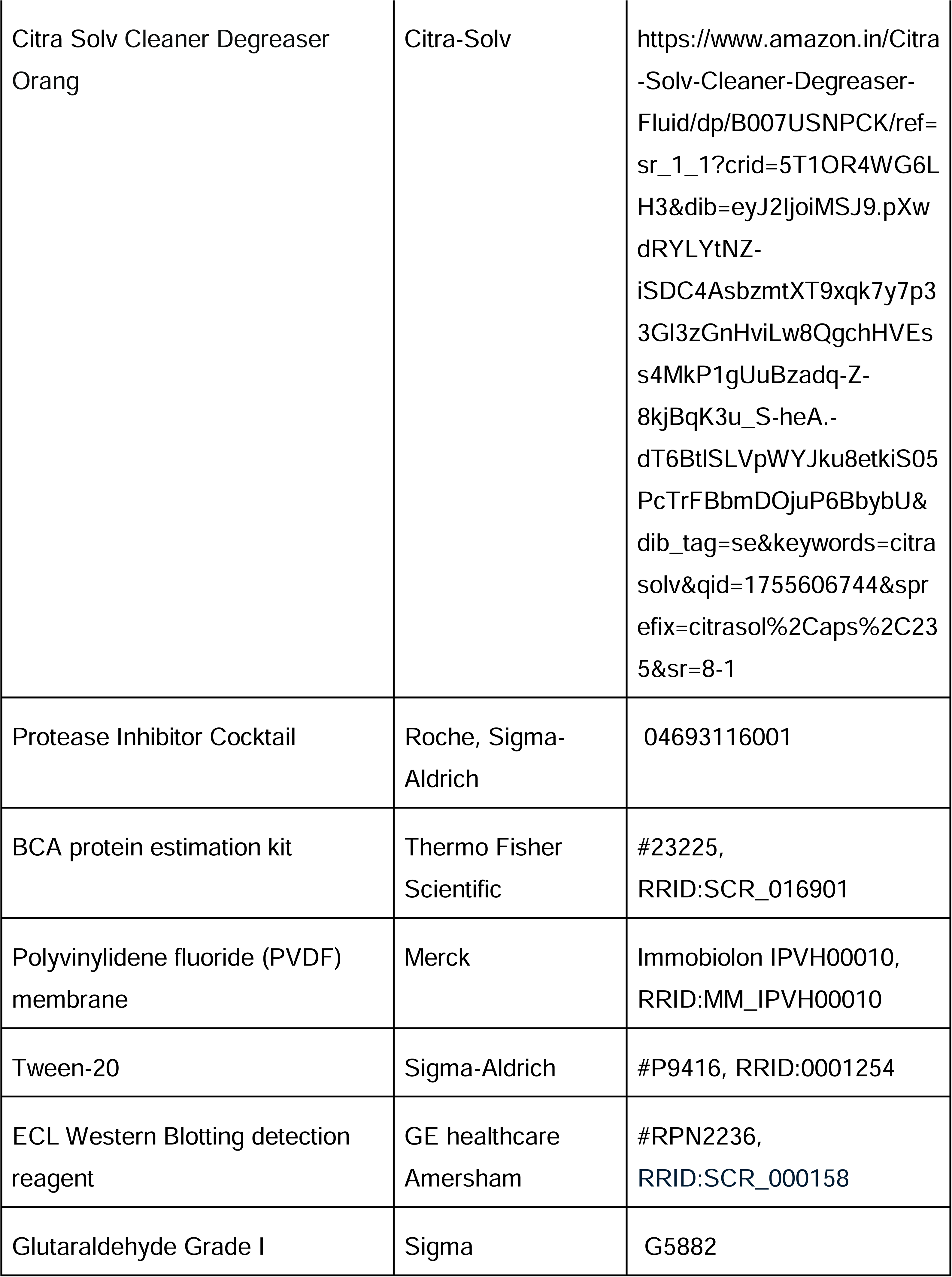
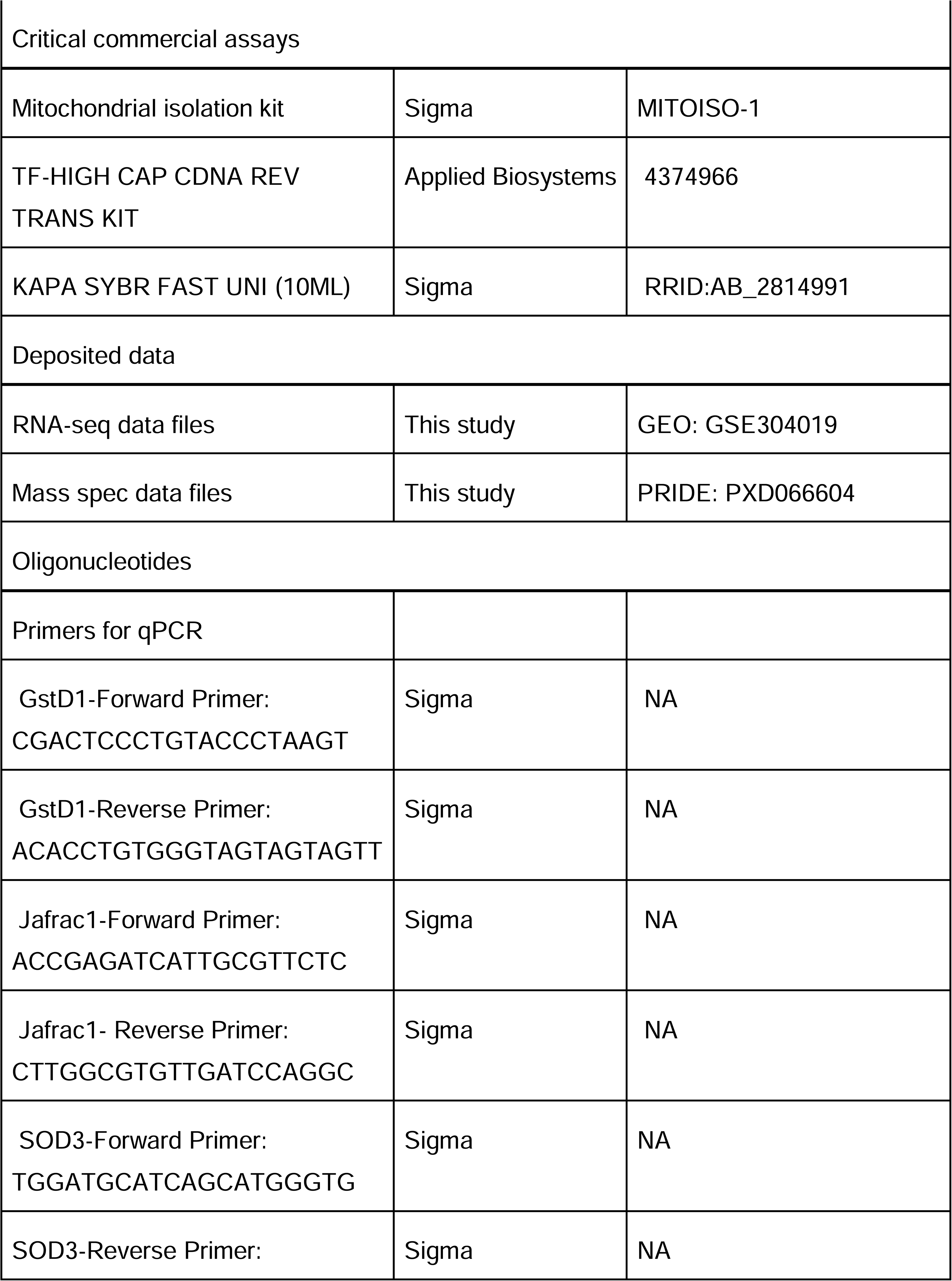
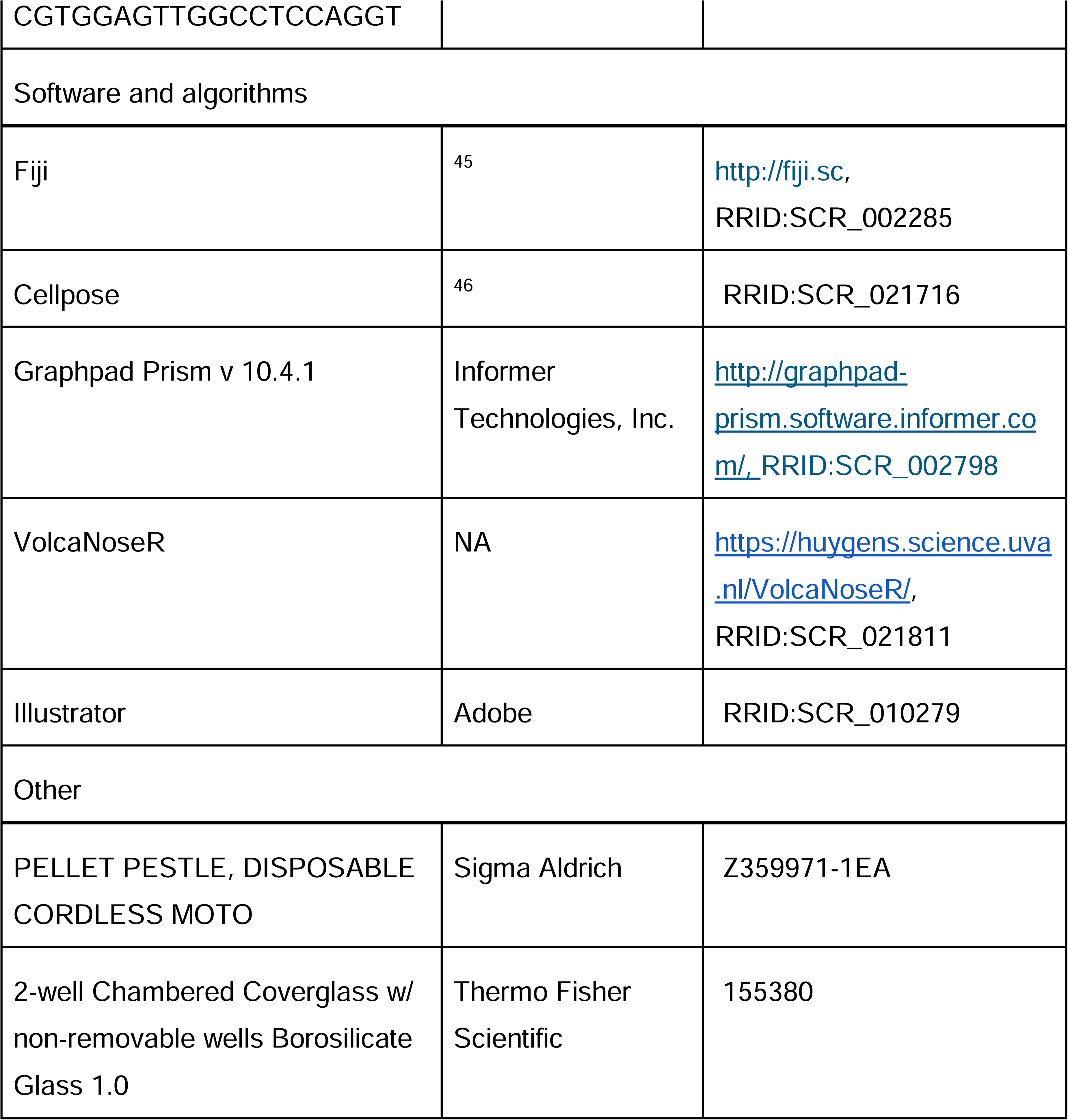

### *Drosophila* stocks and genetics

All the fly crosses were maintained in the standard cornmeal agar medium. dl^i^ (y[1] sc[*] v[1] sev[21]; P{y[+t7.7] v[+t1.8]=TRiP.HMS00727}attP2, BDSC_32934), dl^1^ (dl[1] cn[1] sca[1]/CyO, l(2)DTS100[1], BDSC_3236), dl^4^ (dl[4] pr[1] cn[1] wx[wxt] bw[1]/CyO, BDSC_7096), SOD2^i^ (y[1] sc[*] v[1] sev[21]; P{y[+t7.7] v[+t1.8]=TRiP.HMS00783}attP2, BDSC_32983), opa1^i^ (y[1] sc[*] v[1] sev[21]; P{y[+t7.7] v[+t1.8]=TRiP.HMS00349}attP2, BDSC_32358) were obtained from the Bloomington *Drosophila* Stock Center, Bloomington. *dl*^1^,UASp-Dl^WT^ was obtained from Prof. Girish Ratnaparkhi, IISER Pune, India. UASp-Drp1^SG^ and UASp-Mito-PAGFP were generated in the RR lab. UASp-Mito-GFP was obtained from Rachel Cox (USUHS, Bethesda, MD). SqhGFP67mat/cyo; 3xmScarletCAAX15mat/TM3SbSer was obtained from Prof. Yu-Chiun Wang’s lab. Recombinant fly stocks, nanos-Gal4::UASp-Mito-GFP, nanos-Gal4::UASp-Mito-GFP::sqh-Sqh-mCherry and nanos-Gal4::sqh-Sqh-mCherry were made using standard genetic crosses.

UASp-mito-PAGFP was crossed to *nanos*-Gal4::*sqh*-Sqh-mCherry at 25 °C. *dl*^i^ was crossed to maternal driver line *mat67*;*mat15* carrying maternal 4 *tubulin*-Gal4-VP16, homozygous for chromosome II and III, at 25 °C. *dl*^1^ and *dl*^4^ mutants were crossed to each other at 25 °C. *dl*^1^ mutant was crossed to *nanos*-Gal4::UASp-mito-GFP::*sqh*-Sqh-mCherry at 28 °C. *dl*^1^,UASp-Dl^WT^ was crossed to *nanos*-Gal4::UASp-mito-GFP at 28 °C. For the apical constriction experiment, Sqh-GFP67mat/cyo; 3xmScarlet-CAAX *mat*15/TM3 Sb Ser was crossed to *w*^1118^ and UASp-*Drp1*^SG^ at 25 °C. Embryos were obtained from the F1 generation flies using standard genetic crosses.

### Maintaining the dosage of Gal4 across single and double mutant crosses

Comparable copies of transgenes were maintained across genotypes to ensure comparative dosage and effects of dilution of Gal4 for expression. Double mutant transgenes, *Drp1*^SG^;*opa1^i^* and *Drp1*^SG^;*SOD2*^i^, were crossed with either nanos-Gal4 or *nanos*-Gal4::*sqh*-Sqh-mCherry at 28 °C. Single mutant transgenes, *Drp1*^SG^ and *SOD2*^i^, were crossed to either *nanos*-Gal4::UASp-mito-GFP or *nanos*-Gal4::UASp-mito-GFP::*sqh*-Sqh-mCherry at 28 °C to maintain similar Gal4 amounts as those in the double mutant transgenes.

### Embryonic Lethality Estimation

3-4 hr old embryos were collected from cages. 100 embryos were aligned using a brush on a fresh sugar agar plate in a 10×10 grid and kept at the same temperature as the cage. Unhatched embryos were counted after 24 hrs and 48 hrs. This was done 3 times, and the average of unhatched embryos was calculated and reported as a percentage of the total number of embryos.

### Live imaging

2.5 hr old embryos from cages at 28 °C were dechorionated using bleach for 60-90 seconds, washed and then placed horizontally in a lab-tek chamber. End-on imaging was done to study the process of VF formation. For End-on imaging, after mounting embryos in the lab-tek chamber and adding 1X PBS, each embryo was picked up gently from one end using a single hair brush such that it stands vertically. The lab-tek chamber was then carefully placed at the confocal microscope avoiding any jerk that could cause embryos to fall or tilt. The lab-tek chamber was then screened for completely vertical embryos of the desired stage and imaged using 63X/1.4 NA oil objective on Zeiss laser scanning microscope LSM 710 or Leica laser scanning confocal microscope SP8. The laser power and gain settings were adjusted using the range indicator mode such that the acquired 8-bit image was within the 0-255 range. End-on live imaging was done at 1024×1024 resolution at 2 airy units. 3-5 Z stacks were taken every 30-60 seconds, with 1 μm interval, starting from 45-50 micron depth. For imaging apical constriction, 2.5 hrs old embryos were taken from cages at 25 °C. Embryos at early cellularization were selected and dechorionated using bleach and then washed. Embryos were then placed on an agar slab with the ventral side facing upwards. Lab-tek chamber, with a drop of heptane glue in the centre, was then inverted onto the agar slab gently such that the embryos stuck on their ventral side. 1X PBS was added, and embryos were imaged. 2 color imaging was done using 60X oil objective on the Evident SpinSR10 microscope. The laser power and other acquisition settings were kept the same for control and mutant. 16 bit image was acquired with a time interval of 7.8 seconds.

### Immunostaining

3 hr old embryos were collected from cages at 28 °C for cellularization and gastrulation stages. Embryos were dechorionated using bleach, washed and then fixed using 1:1 Heptane and 8% PFA or 4% PFA (for Drp1 staining) solution for 25 mins. Embryo were then devitellinized either using an insulin needle (hand de-vitellinization) or by shaking in 1:1 Heptane:Methanol (Methanol de-vitellinization for Drp1 staining). After devitellinization, embryos were washed 3 times with 1X PBST (1X PBS with 0.3% Triton X-100) for 5 mins each, blocked using 2% BSA (Himedia, TC545) in 1X PBST for 1 hr at room temperature, then incubated with primary antibodies (anti-Drp1 Booster Bio DZ41314, rabbit, 1:500; anti-Dorsal DSHB 7A4c, mouse, 1:1000; anti-Twist (Eric Wieschaus, Princeton, USA, rabbit, 1:300), anti-Twist (Siegfried Roth, Koln University, Germany, rabbit, 1:500) overnight at 4 °C. Excess primary antibody was removed by giving 3 washes with 1X PBST 5 mins each. Fluorescently labelled dyes (Streptavidin, Alexa Fluor® 488 Conjugate, Invitrogen S32354, 1:1000, Alexa Fluor™ 647 Phalloidin, Invitrogen A22287, 1:1000) and/or secondary antibodies (1:1000), were added and incubated at room temperature for 1 hr after covering with aluminium foil. Following this, 3 washes of 1X PBST were given, and DNA stain Hoechst (Invitogen H3570, 1:1000) was added in the second wash. Finally, embryos were mounted in Slow fade Gold antifade (Thermo Fisher Scientific, S36937) and imaged using Plan Apochromat 63x/1.4 NA, oil immersion objective on Leica laser scanning confocal microscope SP8. Care was taken that the acquired 8-bit image was within the 0-255 range and did not saturate. Imaging was done at 512×512 resolution with a line averaging of 2. 0.5 μm optical sections were acquired for Z stacks at 1AU. For DHE staining, embryos were fixed in 1:1 4% PFA:Heptane for 10 mins and then methanol de-vitellinized. Embryos were taken in 0.1% PBST (1X PBS with 0.1% Triton X-100) and permeabilized with 5% BSA solution made in 1% PBST (1X PBS with 1% Triton X-100) for 90 mins. Blocking was done with 5% BSA made in 0.1% PBST for 30 mins at room temperature. Embryos were incubated in DHE (2 μM in 1X PBS) for 10 mins. Lastly, one wash was given with 1X PBS and then embryos were mounted in Slow fade Gold antifade.

### Photoactivation

Embryos expressing mito-PAGFP and Sqh-mCherry were mounted in end-on orientation. A ring shaped ROI was drawn at the basal region at the end of cellularization/ beginning of gastrulation. The selected ROI was photoactivated using a 405 nm laser at 10% power, using a 60×/1.4 NA Plan apochromat objective on Evident FV4000 inverted confocal microscope. Images were obtained every 30 secs.

### Colchicine treatment via Citrasolv

2-2.5 hr old *nanos*-Gal4::UASp-mito-GFP::*sqh*-Sqh-mCherry embryos were collected from cages at 28 °C. Embryos were treated with bleach for 3 mins and then washed with distilled water. Further, citrasolv treatment was given (1:10 in 1X PBS) for 5 mins on the rocker (embryos were in the sieve). Citrasolv was removed by rinsing the embryos 2-3 times with 1X PBS. Then the embryos were transferred using a brush to a tube containing 1X PBS, and 1mM colchicine (Sigma, #C9754) was added and incubated for 15 mins on the rocker. Finally, the drug was removed and embryos were rinsed with 1X PBS once and then mounted on a lab-tek chamber and imaged using 63X/1.4 NA oil objective on Zeiss laser scanning microscope LSM 710.

### TEM of control gastrulation embryos

A 30 min collection of *yw* embryos at 25 °C was taken and aged till the gastrulation stage. Embryos were then dechorionated with bleach and fixed for 10 min in a mixture of 1 ml glutaraldehyde 25% and 4 ml heptane. Heptane and glutaraldehyde were removed, and embryos were immersed in phosphate-buffered saline (PBS) containing 0.1% Tween 20 and 0.1% bovine serum albumin (BSA). Embryos were hand-devitellinized on a double-stick tape with a hypodermic needle (BD Microlance 3–30Gx1/2″, BD 304000). De-vitellinized embryos of the desired stages were transferred to a freshly prepared fixative solution (2.5% glutaraldehyde in 50 mM cacodylate buffer, pH 7.2–7.4, 0.1% tannic acid) and then stored at 4°C in the same solution overnight. Embryos were post-fixed for 2 h in 1% osmium, 2% glutaraldehyde in 50 mM cacodylate buffer at 4°C. They were then washed in cacodylate buffer, dehydrated in a graded ethanol series, and finally embedded in epon. Ultrathin sagittal sections (80 nm) were prepared with an ultramicrotome, mounted onto copper grids, contrasted with 1% uranyl acetate, and examined with a FEI Tecnai G2 200 kV electron microscope. Images were acquired with a FEI Eagle charge-coupled device (CCD) camera (4096 × 4096 pixels) for 1700× magnification and an Olympus Veleta CCD camera (2048 × 2048 pixels) for 3100× and 8500× magnifications. The procedure was carried out in Institut de Biologie du Développement de Marseille (IBDM) electron microscopy facility (Marseille, France).

### TEM of control and Drp1^SG^ cellularization embryos

1.5 hr embryos were collected, aged for 2 hrs, dechorionated using bleach for 1 min and then rinsed with distilled water. Embryos were then transferred to a scintillation vial containing a 1:1 solution of 3% glutaraldehyde (Grade I, Sigma, G5882) and heptane for 2 hr at 4°C on the rocker for fixation. After fixation, all the solution was removed, fresh heptane was added and then embryos were hand-devitellinized using an insulin needle. Devitellinized embryos were collected in a 0.5ml eppendorf tube. Two washes were given with 0.1M sodium cacodylate buffer for 10 mins each and post-fixed in 1 % Osmium tetroxide (OsO4) for 1 hr at 4 °C. The tissues were en-block stained in 2% uranyl acetate followed by dehydration in an ascending alcohol series for 10 minutes each. The tissues were infiltrated in Araldite resin (A and B) and finally embedded in Araldite resin B and polymerized at 60 °C for 48 hrs. Ultrathin sections of 70 nm were cut using an ultramicrotome (LeicaUC7) and collected on copper slot grids. The sections were contrasted with lead citrate. The images were acquired using a Transmission Electron Microscope, JEM 1400 Plus, JEOL (Japan) at 120 kV. The sample embedding, sectioning and imaging were carried out in the Transmission Electron Microscope facility, ACTREC, Tata Memorial Centre, Kharghar, Navi Mumbai, India.

### Western Blotting

3 hr old embryos were collected from cages at 28 °C and dechorionated using bleach for 1 min, followed by washing in distilled water. Then embryos were crushed on ice in 150-200 μl lysis buffer (1%Triton-X 100, 50 mM Tris HCl; pH 8.0, 150 mM NaCl, Protease Inhibitor Cocktail; PIC 1:50) using a homogeniser gun. The homogenate was then centrifuged at 15000 g for 30 mins at 4 °C. The supernatant was removed carefully, avoiding the lipid layer at the interface, and stored at −80 °C. Protein concentration of the supernatant was determined using BCA protein estimation kit (Thermo Fisher Scientific #23225). A sample for SDS-PAGE was prepared by boiling the protein sample (supernatant) in 1X Laemmli buffer at 95 °C for 10 mins. The denatured protein sample was then loaded into 10% or 12% polyacrylamide gel followed by transfer onto a 0.45μm polyvinylidene fluoride (PVDF) membrane (Merck Immobiolon IPVH00010). Post transfer, the blot was blocked for 1 hr at room temperature with 5% skimmed milk or 5% BSA dissolved in 1X TRIS buffer saline (TBS) with 0.1% Tween-20 (Sigma-Aldrich, #P9416) detergent. After blocking, the primary antibodies (Drp1 1:5000; β-actin Cell signaling #4967, rabbit, 1:1000; ATP beta abcam ab14730, mouse, 1:2000) was added and kept at 4 °C for 12-16 hrs. Following this, the blot was then washed, 3 washes 15 mins each, using 1X TRIS buffer saline (TBS) with 0.1% Tween-20. Then, secondary antibody is added for 1 hr at a concentration of 1:10,000 and kept at room temperature. Blot is then again washed thrice and then developed using ECL Western Blotting detection reagent (GE Healthcare Amersham, #RPN2236). For quantitative analysis, the image was first inverted. Background subtraction was done. The intensity of bands was normalized by dividing it by the intensity of the loading control.

### Mitochondria Isolation

Mitochondria are isolated from embryos using the mitochondria isolation kit (Sigma, MITOISO-1). 3 hr old embryos are taken from cages at 28 °C and dechorionated using bleach for 1 min. Dechorionated embryos are rinsed with a solution EB-A twice. Then, embryos are crushed on ice in 200 μl EB-A with 2mg/ml BSA and centrifuged at 600g for 5 mins at 4 °C. The pellet is discarded, and the supernatant is centrifuged at 11,000g for 10 mins at 4 °C. The supernatant was discarded. The pellet is then resuspended in 100 μl EB A buffer and centrifuged at 600 g for 5 mins at 4 °C. Following this, the pellet obtained is discarded as it contains the nuclear fraction. The supernatant is centrifuged at 11,000 g for 10 mins at 4 °C. The pellet obtained after this spin is enriched in mitochondria, and it is resuspended in a storage buffer and stored at −20 °C.

### RNA sequencing

For sample preparation for RNA sequencing, embryos were manually sorted to avoid non-developing embryos and to select for only cellularizing and gastrulating embryos. Firstly, egg laying was synchronised by discarding the first 2 hr collection. For genotypes that laid enough embryos, 1 hr collection was taken and aged for 1.5-2 hrs, whereas for mutants that did not lay enough embryos 2.5-3 hr collection was taken directly after synchronisation from cages at 28 °C. Embryos were dechorionated using bleach and washed with distilled water. Then, around 50-60 embryos were taken, using a brush, on a slide, and around 20-30 μl 1X PBS was added on top of the embryos. Embryos of desired stages were then sorted using a single-haired brush under a stereo microscope. These embryos were taken in a tube and crushed in TriZol (Qiagen) using a homogeniser gun, snap frozen in liquid nitrogen, and then stored at −80 °C. Multiple embryo collections were pooled such that there were 70-100 embryos (equal proportion of cellularizing and gastrulating embryos) in each replicate. RNA was isolated using the TRIZOL method. The concentration of RNA was measured using Qubit RNA BR Assay (Invitrogen, Cat# Q10211). RNA purity was checked using QIAxpert, and RNA integrity was checked on TapeStation using RNA ScreenTapes (Agilent, Cat# 50675579). The samples were taken for the library preparation and sequencing.

#### Library Prep Protocol

KAPA RNA HyperPrep Kit (cat #0000141759) protocol was used to prepare libraries for mRNA sequencing. An initial Concentration of 200ng of total RNA was taken for the assay. mRNA molecules are captured using magnetic oligo Poly(T) beads (KAPA mRNA capture kit, Cat# 7962240001). Following purification, the enriched mRNA was fragmented using divalent cations under elevated temperatures (94 degrees for 6 minutes). The cleaved RNA fragments were copied into first-strand cDNA using reverse transcriptase and random priming. Combined 2nd strand synthesis and A-tailing is done, which converts the cDNA:RNA hybrid to double-stranded cDNA (dscDNA), incorporates dUTP into the second cDNA strand, and adds dAMP to the 3’ ends of the resulting dscDNA. This is followed by Adapter Ligation, in which dsDNA indexing adapters with 3’ dTMP overhangs are ligated to library insert fragments. The adapter-ligated products were then purified and enriched carrying appropriate adapter sequences at both ends using high-fidelity, low-bias PCR (the strand marked with dUTP is not amplified, allowing strand-specific sequencing) using the following thermal conditions: initial denaturation 98 °C for 45sec; 13 cycles of − 98 °C for 15sec, 60 °C for 30sec, 72 °C for 30sec; final extension of 72 °C for 1mins. PCR products are then purified and checked for fragment size distribution on Fragment Analyzer using HS NGS Fragment Kit (1-6000bp) (Agilent, Cat# DNF-474-1000).

#### Sequencing Protocol

Prepared libraries were quantified using Qubit HS Assay (Invitrogen, Cat# Q32854). The obtained libraries were pooled and diluted to the final optimal loading concentration. The pooled libraries were then loaded onto Illumina NovaSeq V1.5 instrument at Medgenome, India, to generate 40M, 150bp paired-end reads/sample.

#### Read quality check

The following parameters were checked from fastq file - Base quality score distribution, Sequence quality score distribution, Average base content per read, GC distribution in the reads, PCR amplification issue, Check for over-represented sequences and Adapter trimming. Based on the quality report of fastq files, sequence reads were trimmed where necessary to only retain high quality sequence for further analysis. In addition, the low-quality sequence reads were excluded from the analysis. The adapter trimming was performed using Trimmomatic (v-0.36).

#### Contamination removal

For the RNA-Seq analysis, the unwanted sequences were removed, especially nonpolyA tailed RNAs from the sample (assuming that poly-A tailed RNAs are sequenced). The unwanted sequences include - mitochondrial genome sequences, ribosomal RNAs, transfer RNAs, adapter sequences and others. Contamination removal was performed using Bowtie2 (2.2.4).

#### Read alignment

The paired-end reads were aligned to the reference *Drosophila melanogaster* genome using HISAT2 (2.1.0) to quantify reads mapped to each transcript. Alignment percentage of reads was in the range of 91-95 percent for all the samples.

#### Expression estimation

The aligned reads were used for estimating the expression of the genes. The raw read counts were estimated using FeatureCount (1.5.2).

#### Differential expression analysis

The raw read counts were normalized using DESeq2. The ratio of normalized read counts for treated over control was taken as the fold change. Genes were first filtered based on the p-value (≤ 0.05). A distribution of these log2 (foldchange) values were found to be normally distributed. Those genes which were found to have -1≤ log2(foldchange) ≥1 were considered statistically significant. Gene ontology of the significantly differentially expressed gene set was performed using ShinyGO 0.81. Volcano plot was plotted using VolcaNoseR. Heat maps were plotted in GraphPad Prism 10.4.1 using log (CPM+1) values for genes which were unchanged and z-normalized values for differentially expressed genes. The RNA isolation, sequencing and analysis was done with MedGenome Labs, Bangalore, India.

The RNA sequencing data from this study are available at the NCBI GEO database under the accession number GSE304019.

### RT-PCR

The concentration of RNA was adjusted to 1μg using nuclease-free water. High-capacity cDNA reverse transcription kit (Applied Biosystems, 4374966) was used for the synthesis of cDNA using the manufacturer’s protocol. cDNA was used at a dilution of 1:5 for all reactions. Primers (Sigma) used were the following:

GstD1-Forward Primer: CGACTCCCTGTACCCTAAGT

GstD1-Reverse Primer: ACACCTGTGGGTAGTAGTAGTT

Jafrac1-Forward Primer: ACCGAGATCATTGCGTTCTC

Jafrac1- Reverse Primer: CTTGGCGTGTTGATCCAGGC

SOD3-Forward Primer: TGGATGCATCAGCATGGGTG

SOD3-Reverse Primer: CGTGGAGTTGGCCTCCAGGT

The *ΔΔCt* method was used to calculate the gene expression changes among comparative genotypes (control and Drp1^SG^)

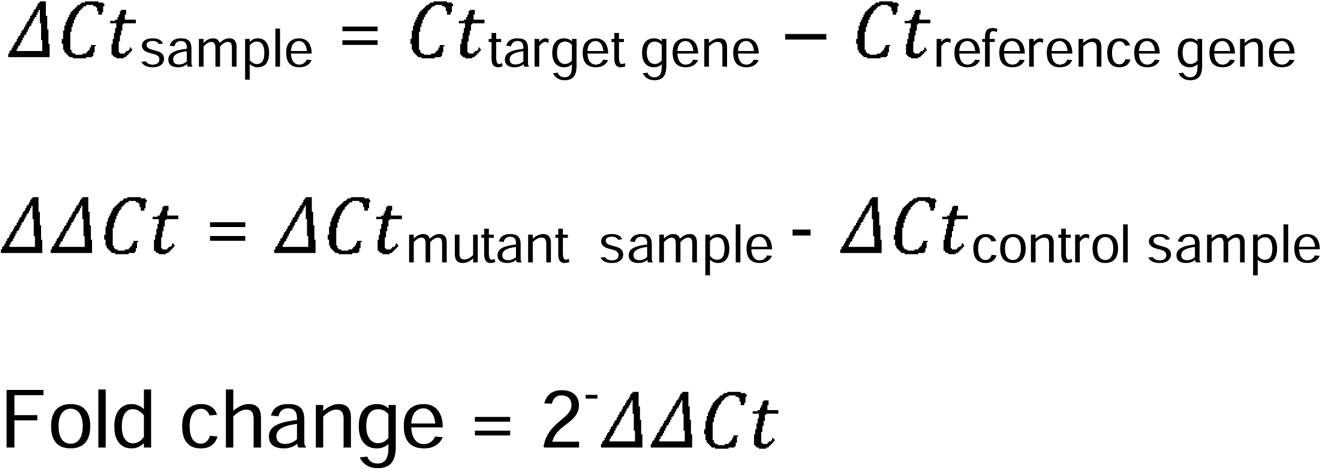

Where, Ct is the cycle threshold or cq value. The reference gene is the housekeeping gene *rp49*. Comparisons between genotypes were made, and fold change was plotted in the form of a bar chart to represent the difference in levels of gene expression.

### Mass Spectrometry

Two kinds of samples were prepared for mass spectrometry, total protein sample and mitochondrial protein sample. Total protein sample was prepared similarly to the sample prep for western blots, wherein embryo lysate in lysis buffer is centrifuged at 15000 g for 30 mins at 4 °C and the supernatant is used for mass spec. For the mitochondrial protein sample, mitochondria were isolated from embryos as mentioned above. The final mitochondrial pellet was resuspended in a buffer containing 5% SDS and 0.1M Tris pH 8.0, and used for mass spec. The following processes were done with Vproteomics, New Delhi, India.

#### Sample Preparation for Mass Spec

25 μg of the protein sample was first reduced with 5 mM TCEP and subsequently alkylated with 50 mM iodoacetamide. The protein was then digested with Trypsin at a 1:50 Trypsin-to-lysate ratio for 16 hours at 37 °C. After digestion, the mixture was purified using a C18 silica cartridge and then concentrated by drying in a speed vac. The resulting dried pellet was resuspended in buffer A, which consists of 2% acetonitrile and 0.1% formic acid.

#### Mass Spectrometric Analysis of Peptide Mixtures

For mass spectrometric analysis, all the experiments were performed on an Easy-nLC-1000 system (Thermo Fisher Scientific) coupled with an Orbitrap Exploris 240 mass spectrometer (Thermo Fisher Scientific) and equipped with a nano electrospray ion source. 1 μg of phosphopeptides sample was dissolved in buffer A containing 2% acetonitrile/0.1% formic acid and resolved using Picofrit column (1.8-micron resin, 15cm length). Gradient elution was performed with a 0–38% gradient of buffer B (80% acetonitrile, 0.1% formic acid) at a flow rate of 500nl/min for 96 mins, followed by 90% of buffer B for 11 min and finally column equilibration for 3 minutes. Orbitrap Exploris 240 was used to acquire MS spectra under the following conditions: Max IT = 60ms, AGC target = 300%; RF Lens = 70%; R = 60K, mass range = 375−1500. MS2 data was collected using the following conditions: Max IT= 60ms, R= 15K, AGC target 100%. Lock mass option was enabled for polydimethylcyclosiloxane (PCM) ions (m/z = 445.120025) for internal recalibration during the run. MS/MS data was acquired using a data-dependent top20 method dynamically choosing the most abundant precursor ions from the survey scan, wherein dynamic exclusion was employed for 30s.

#### Data Processing

RAW files generated were analyzed with Proteome Discoverer (v2.5) against the Uniprot *Drosophila melanogaster* database. For dual Sequest and Amanda search, the precursor and fragment mass tolerances were set at 10 ppm and 0.02 Da, respectively. The protease used to generate peptides, i.e. enzyme specificity, was set for trypsin/P (cleavage at the C terminus of “K/R: unless followed by “P”). Carbamidomethyl on cysteine as fixed modification and oxidation of methionine and N-terminal acetylation were considered as variable modifications for database search. Both peptide spectrum match and protein false discovery rate were set to 0.01 FDR.

Further downstream analysis was performed using in-house R-scripts. For differential expression analysis limma package in R was used. For differential protein abundance analysis, the abundance values were first Log2 transformed. Then the data was filtered based on valid values in at least 70% of the replicates within each condition. Missing values were imputated based on normal distribution. Abundance values were median-based quantile normalised. Bioreplicates were categorized and grouped in Control and Treated groups. Post-processing, t-test was applied which provided the pValues and log2 fold change values. Proteins having p value≤0.05 were considered significant. log2 fold change ≥0.3 and ≤-0.3 was considered significant. Abundance values of significant proteins were Z-scale transformed. Heatmap was generated on these proteins Z-normalized data using GraphPad Prism 10.4.1. Gene ontology of the significantly differentially expressed proteins was performed using ShinyGO 0.81. Volcano plot was plotted using VolcaNoseR. All the raw and processed mass spectrometry data can be accessed at the PRIDE repository with the identifier PXD066604.

## Image Analysis and quantification

### Mitochondrial fluorescence in the ventral furrow

Mean mitochondrial fluorescence was measured in apical (above the nucleus) and basal (below the nucleus) regions of cells at 3 different regions of the embryo: Ventral (cells undergoing constriction, marked in red), neighbouring (cells adjacent to the constricting cells, marked in green), and lateral (cells on the lateral region of the embryo, marked in blue). 5 different time points were chosen, based on the shape and extent of invagination of the furrow being formed, starting from the end of cellularization till the time point when the furrow closes. The background intensity was taken from a region within the embryo that did not contain any visible fluorescence and subtracted from the apical and basal intensities. Ratio of background subtracted apical:basal intensities were normalized with the ratio of the first time point i.e., end of cellularization and plotted as Mean ± SEM using GraphPad Prism 10.4.1. Image analysis was done using ImageJ.

### Quantification of mitochondria in TEM

Mitochondria in the EMs were identified based on structure and electron density. Mitochondria in each cell were counted manually, and their size and circularity were measured using ImageJ.

### Mitochondrial and Myosin II fluorescence in photoactivation

Fluorescence of mitochondria and Myosin II was followed in the apical and basal regions of the ventral and lateral cells. Background normalization was done by subtracting the mean intensity of a region within the embryo. Background subtracted mean apical:basal intensities of mitochondria and Myosin II for the 2 regions (ventral and lateral) were measured across time and plotted as Mean ± SEM using GraphPad Prism 10.4.1.

### Measurement of mitochondrial area in apical and basal sections

Single planes at the apical and basal regions were used for analysis. Firstly, background normalization was done by subtracting the mean intensity value. The thresholding was done, and particles greater than 0.05 μm^2^ were marked using the “particle analyser” tool in ImageJ. Mean particle size was determined for each embryo. The data were plotted as a violin plot and analyzed with a Mann–Whitney test using GraphPad Prism 10.4.1.

### Measurement of relative mitochondrial area in apical sections

Apical stacks of late cellularization embryos were selected using phalloidin and Hoechst as a reference. Particles greater than 0.05 μm^2^ were selected using thresholding. Relative mitochondrial area was obtained by dividing the total area of all thresholded mitochondrial particles in the apical stacks by the total area of the imaging field and expressed as a percentage. The data were plotted and analyzed using GraphPad Prism 10.4.1.

### Dorsal and Twist intensity

Nuclei were marked using Hoechst staining. Fluorescence intensity of Dorsal and Twist was measured in the marked nuclei. Mean ventral to dorsal fluorescence intensity was plotted and compared using the Mann-Whitney test in GraphPad Prism 10.4.1.

### Mitochondrial fluorescence in Dorsal overexpression

ROI was drawn, using the segmented line tool, in the apical region of the entire embryo (end-on) in control and *dl*^1^,UASp-Dorsal and the mean mitochondrial fluorescence was measured. The perimeter of the embryos was measured and normalized to arbitrary units in percentage such that the ventral midpoint is marked as 0, left to that is 0 to -50, and right to that is 0 to 50. Mean of the normalized intensities at every 1.0 airy unit was plotted using GraphPad Prism 10.4.1.

### Quantification of apical area and Myosin II intensity during apical constriction

For representation and quantification, a single Z-plane at 5 μm depth from the apical cortex was taken for ScarletCAAX. For Myosin II, average intensity Z-projections were taken of the planes that showed Sqh-GFP signal. The time point when the furrow reaches a depth of 2.5-2.7 μm was taken as the end, and 7.8 mins before that was taken as the first time point. Cells were segmented and tracked, and their area and Sqh-GFP intensity were measured using Cell Pose and Trackmate. Segmented cells were color-coded based on the area. Area and Myosin II intensity of 40 cells from one control and mutant embryo were plotted as a heat map in GraphPad Prism 10.4.1. Mean apical area and mean Myosin II intensity were normalized to their respective first time and plotted as Mean ± SEM using GraphPad Prism 10.4.1.

### Quantification of DHE fluorescence

The mean fluorescence intensity of DHE was calculated for a single plane located just above the nucleus. The mean intensity values of all genotypes were normalized with the average mean intensity of the control, and plotted using GraphPad Prism 10.4.1 and compared using the Mann-Whitney test.

### Measurement of ventral furrow width

A single time point was chosen based on the furrow shape (when the furrow was U-shaped- T4). The furrow width was calculated by drawing a line across the furrow at the point where the Myosin II signal (Sqh-mCherry) ended. The mean values were plotted as a violin plot in GraphPad Prism 10.4.1 and compared using the Mann-Whitney test.

### Myosin II fluorescence in the ventral furrow

Mean Myosin II (Sqh-mCherry) intensity was measured in the ventral cells by drawing an ROI around the apical Myosin II signal. Background intensity was measured from a region within the embryo, which did not have visible fluorescence. Background subtracted mean Myosin II intensity was normalized with the first time point and plotted with SEM using GraphPad Prism 10.4.1.

### Statistics and n values

The Mann-Whitney test was applied for statistical analysis. n values ranged from 3 to 27 and are mentioned in the respective figure legends for each experiment. Significance levels are indicated as follows: p≥0.05 (ns), p<0.05 (*), p<0.01 (**), p<0.001 (***), p<0.0001 (****).

## Results

### Mitochondria are enriched apically in the ventral furrow cells during gastrulation

Mitochondria are fragmented, dispersed and highly abundant in the early *Drosophila* embryos (Chowdhary et al., 2020, 2017). We imaged mitochondria during cellularization and gastrulation with fluorescently coupled streptavidin (Chowdhary et al., 2017). As seen previously, mitochondria were enriched apically as cellularization proceeded, with mitochondria present above the nucleus in late stages of cellularization (Figure 1B). We further evaluated the distribution of mitochondria in gastrulation. Mitochondria appeared as small, punctate structures predominantly in apical regions, above the nucleus, in VF cells as compared to the basal regions (Figure 1C). We performed live imaging of mito-GFP expressing embryos during gastrulation in an end-on orientation to study the dynamics of mitochondria. The end-on orientation enabled the visualization of apico-basal mitochondrial dynamics in the dorsal, lateral, and ventral axes of each embryo simultaneously. We observed an apical enrichment of mitochondria in constricting VF cells compared to the neighboring and the lateral cells. This enrichment increased as gastrulation proceeded and peaked when cells invaginated to form the VF (Figure 1D, Video 1). We measured the mean apical and basal mitochondrial fluorescence in 3 regions, constricting ventral cells (VF cells, red), immediately neighbouring cells (green) and lateral cells (blue) across 5 time points of VF formation. The apical:basal mean mitochondrial fluorescence ratio was normalised to the first time point at the end of cellularization. This quantification showed that the apical mitochondrial intensity increased with time in the ventral cells but remained almost constant in the neighbouring and lateral cells (Figure 1E). We observed a similar abundance of mitochondria in the apical regions of the VF cells compared to the lateral cells in transmission electron micrographs (Figure 1F). The numbers of apical mitochondria per cell were significantly increased in ventral cells compared to lateral cells (Figure 1G). In summary, we found that small, punctate mitochondria were enriched in apical regions in the VF cells during gastrulation.

### Mitochondria translocate apically in the ventral furrow cells in a microtubule-dependent manner

Mitochondria translocate in cells with the help of motor proteins in a microtubule-dependent manner from the perinuclear region to the periphery in migrating cells and from the cell body to the synapse in neurons and from basal to apical regions in cells in *Drosophila* cellularization (Chowdhary et al., 2020; Desai et al., 2013; Pilling et al., 2006; Zhao et al., 2013). We assessed whether apical migration in VF cells occurred from basal regions and if this was dependent on microtubules. Live embryos expressing matrix-labelled photoactivatable mito-GFP (mito-PAGFP) and Sqh-mCherry were imaged in the end-on orientation. A region of interest in the basal section of all cells was photoactivated at the end of cellularization or the beginning of gastrulation (Figure 2A, white ROI). Remarkably, we saw an increased movement of mitochondria from the basal to the apical regions in the VF cells as compared to lateral cells immediately after photoactivation (Figure 2A, Video 2). The mitochondrial fluorescence in apical regions of ventral cells increased to a significantly greater extent as compared to lateral cells (Figure 2B). This demonstrated that mitochondrial migration from the basal region to the apical regions was enhanced explicitly in the ventral cells during gastrulation. The increase in mitochondrial intensity was accompanied by an increase in the Myosin II fluorescence in the ventral cells (Figure 2A and B). We observed mitochondria migrate linearly in these cells (Figure 2A, Zoomed in, white arrowheads). To further assess whether the migration was microtubule-dependent, we treated embryos expressing mito-GFP with the microtubule-depolymerizing drug colchicine at the end of cellularization and imaged them live in an end-on orientation during the process of gastrulation. Colchicine treated embryos showed reduced VF invagination, as has been reported in previous studies (Ko et al., 2019). Apical mitochondrial enrichment in the constricting cells was significantly reduced, and mitochondria were instead enriched at the basal side (Figure 2C). We quantified the mean apical:basal mitochondrial intensity in the constricting cells at a time point showing comparable constriction in the ventral cells in control and colchicine treated embryos, and found that it was significantly reduced in colchicine treated embryos compared to the control embryos (Figure 2D). These data confirmed that mitochondria migrate in a microtubule-dependent manner from the basal to the apical side in VF cells during gastrulation.

**Figure 2.**
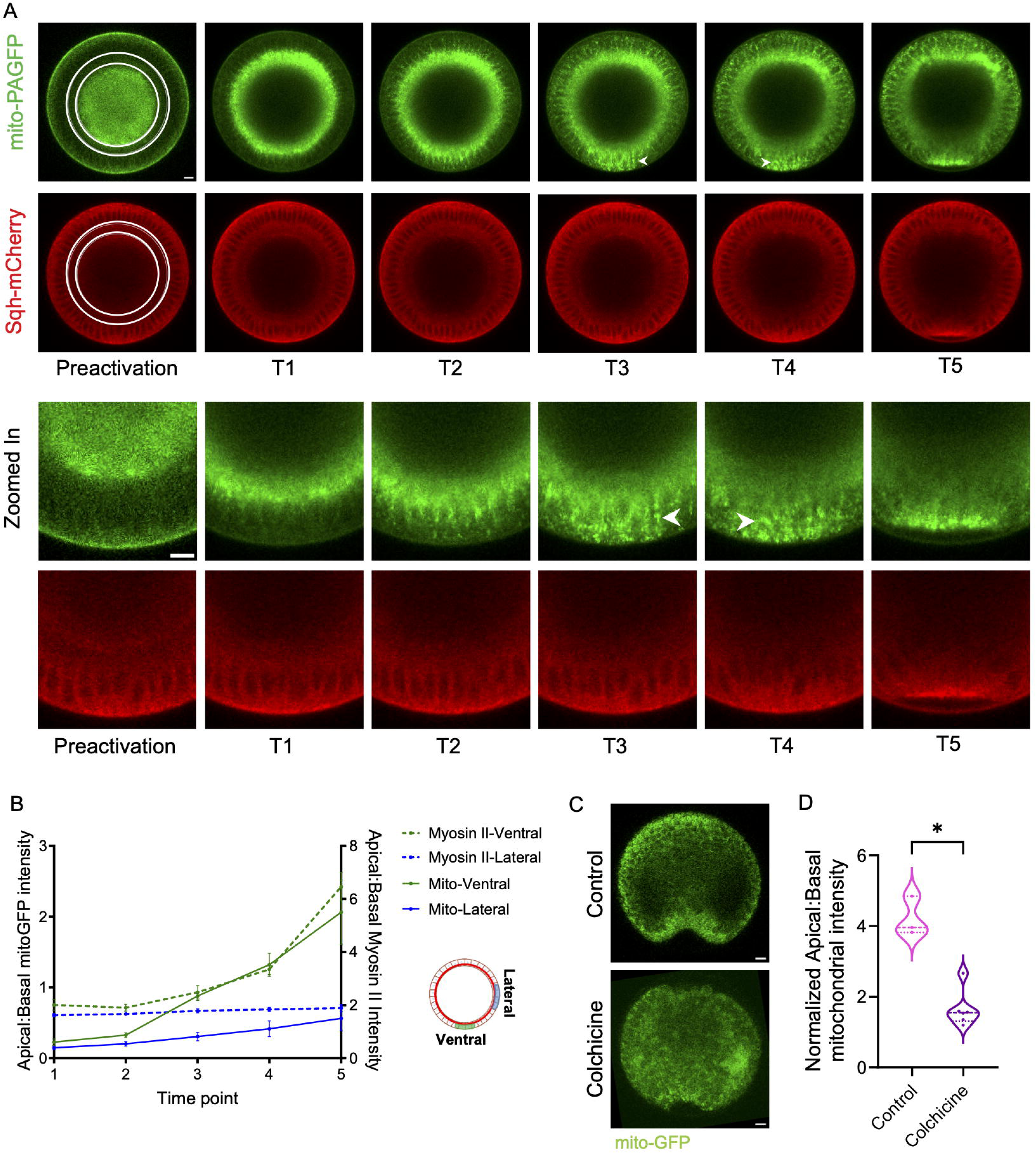
Mitochondria translocate apically from basal sections in the ventral furrow cells in a microtubule-dependent manner. (A) Representative embryo expressing mito-PAGFP and Sqh-mCherry was photoactivated in the white ROI at the end of cellularization or the beginning of gastrulation. Representative snapshots are shown from the movie before and after photoactivation (T1-T5). Zoomed in images show mitochondria being translocated from basal to apical regions in ventral cells on linear tracks. (B) The plot shows mean apical:basal fluorescence intensity of mitochondria and Myosin II measured in ventral and lateral cells (n=3), as shown in the schematic. Data is represented as Mean ± SEM. (C) Representative images of mito-GFP expressing embryos treated with colchicine (n=6) and the respective control (n=3). Colchicine treated embryos show reduced mitochondrial migration in the ventral cells and clustering of mitochondria in the basal regions (yellow arrowhead). (D) The plot shows mean mito-GFP fluorescence intensity of the ventral cells expressed as a ratio of apical to basal regions. The violin plot shows median and quartiles. The statistics were performed by two-tailed unpaired Mann-Whitney test, *p<0.05. Scale Bar: 10μm (A,C).

### Dorsal/NFkB regulates apical translocation of mitochondria in gastrulation

Dorso-ventral patterning in *Drosophila* embryos is activated by the Dorsal/NFkB (Dorsal) signaling pathway in ventral cells (Belvin and Anderson, 1996). We further assessed the role of the Dorsal pathway in regulating the migration of mitochondria in the VF cells. We used the *dl*RNAi and transheterozygous Dorsal null mutants (*dl*^4^/*dl*^1^) to assess the mitochondrial distribution in these embryos. In both cases, embryos showed 100% lethality and did not proceed beyond cellularization to the formation of the VF. We therefore assessed the mitochondrial dynamics in cellularization in Dorsal depleted embryos. *dl*RNAi and *dl*^4^/*dl*^1^ mutants were stained with streptavidin, and mitochondrial morphology and distribution were assessed. We found that the apical mitochondrial fluorescence in *dl*RNAi and *dl*^4^/*dl*^1^ embryos was reduced compared to the controls (Figure 3A). The percentage of apical area occupied by mitochondria in late cellularizing embryos was calculated using a threshold for mitochondrial fluorescence. The area occupied by mitochondria in apical sections in *dl*RNAi and *dl*^4^/*dl*^1^ embryos was significantly reduced as compared to control embryos (Figure 3B). However, the mitochondrial size in the apical and basal sections was similar in *dl*RNAi, *dl*^4^/*dl*^1^ and control embryos (Figure 3C). Loss of Dorsal decreased the apical mitochondrial migration in cellularization but did not change the mitochondrial morphology. We further assessed haplo-insufficient heterozygous *dl*^1^/+ embryos for defects in mitochondrial migration to the apical side. *dl*^1^/+ embryos showed 74% lethality and have been previously shown to cause ventral patterning defects due to haploinsufficiency at higher temperatures (Hegde et al., 2022; Isoda et al., 1992; Nüsslein-Volhard et al., 1980; Simpson, 1983). First, we assessed the mitochondrial migration in cellularization in *dl*^1^/+ embryos. The relative apical area occupied by mitochondria was significantly lower than that of the controls. The mitochondrial size in apical and basal sections was comparable to that of control embryos (Figures 3A-C). Further, we imaged mitochondria in live *dl*^1^/+ embryos expressing mito-GFP mounted in the end-on orientation (Video 3). 24% (4/17) *dl*^1^/+ embryos did not form the VF, 47% (8/17) of embryos showed a shallow defective furrow (Figure 3D, white arrowhead), and 29% (5/17) of embryos showed VF formation similar to controls. In the *dl*^1^/+ embryos with defective furrow, the apical to basal mitochondrial fluorescence in the ventral cells was significantly lower compared to controls (Figure 3E).

**Figure 3.**
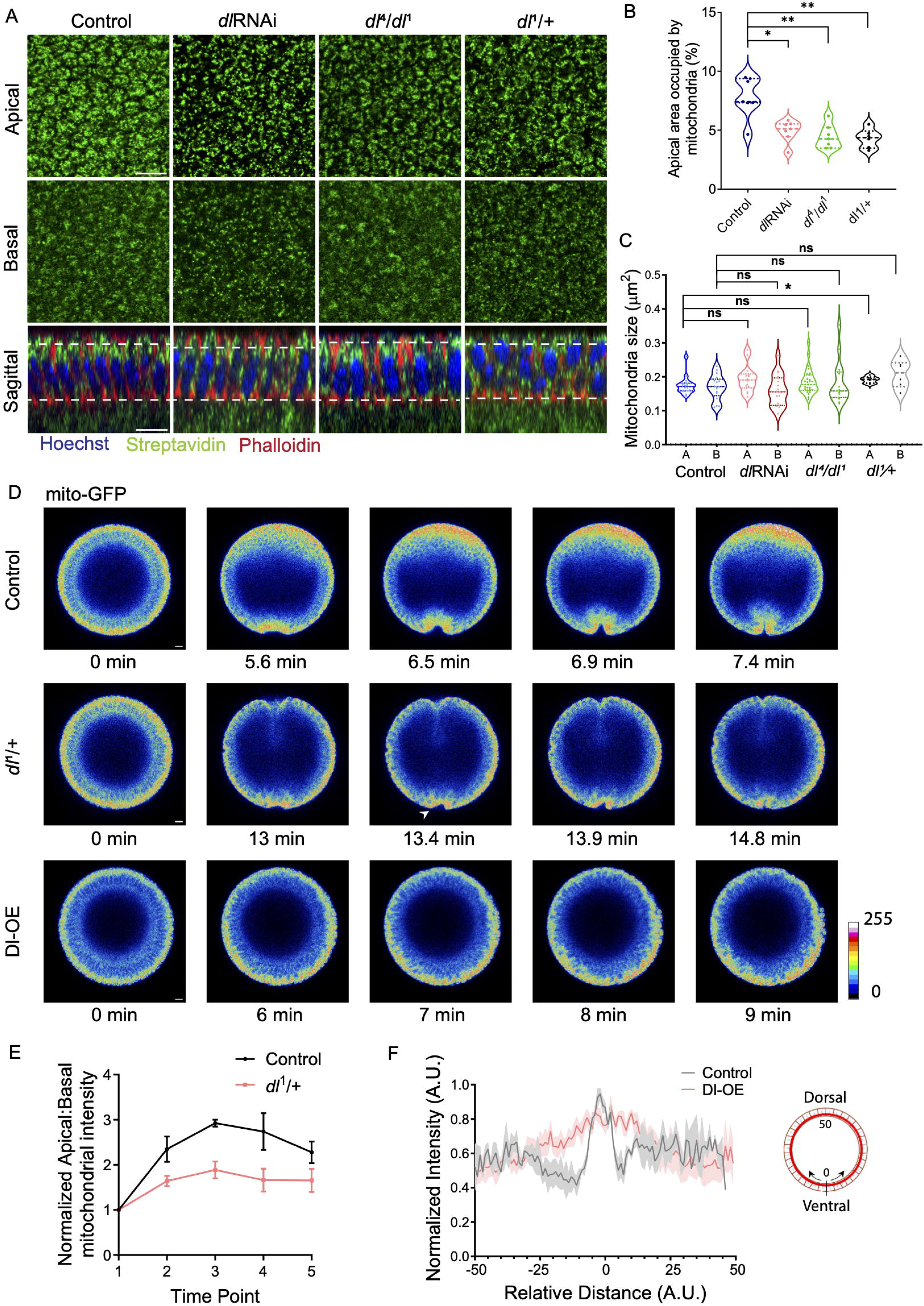
Dorsal regulates apical translocation of mitochondria. (A-C) Dorsal regulates apical translocation of mitochondria during cellularization. (A) Representative optical planes from grazing sections of apical, basal regions and sagittal sections of control, *dl*RNAi, *dl*^4^/*dl*^1^ and *dl*^1^/+ embryos stained for mitochondria (fluorescent streptavidin, green) and DNA (Hoechst, blue) are shown. (B) The plot shows quantification of the relative area occupied by mitochondria in apical sections from *dl*RNAi (n=7), *dl*^4^/*dl*^1^ (n=7) and *dl*^1^/+ embryos (n=6) and control (n=7) embryos. (C) The plot shows mean mitochondrial size quantified from a single optical plane in the apical (A) and basal (B) section. Number of embryos in control apical n=16, basal n=17; *dl*RNAi apical n=14, basal n=16; *dl*^4^/*dl*^1^ apical n=27, basal n=15 and *dl*^1^/+ apical n=6, basal n=6. (D) Representative snapshots from live movies of gastrulating control, *dl^1^*/+ and Dorsal overexpression (Dl-OE) embryos expressing mito-GFP are shown in the 16 color rainbow scale on an 8 bit image with black as 0 and white as 255. (E) The plot shows mean fluorescence intensity from apical sections as a ratio to basal sections at the VF in *dl^1^*/+ (n=4) and control (n=3) embryos. Data is represented as Mean ± SEM. (F) The plot shows mean mitochondrial intensity from mito-GFP at apical regions starting from the ventral midpoint (0) to the dorsal side in controls (n=3) and Dl-OE (n=6). Data is represented as Mean ± SEM. The violin plots show median and quartiles. The statistics were performed by two-tailed unpaired Mann-Whitney test, ns p≥0.05, *p<0.05, **p<0.01. Scale Bar: 10μm (A,D).

We further tested the mitochondrial migration in embryos overexpressing Dorsal. We overexpressed Dorsal in the background of *dl*^1^ (*dl*^1^,UASp-Dorsal) using *nanos*-Gal4, UASp-mito-GFP (referred to as Dl-OE). Dorsal overexpression led to localization of Dorsal and Twist antibodies on both the dorsal and ventral side in 88% of the embryos in contrast to controls showing an expression on the ventral side (Figure S1A). This increase in expression led to a decrease in the ratio of ventral:dorsal nuclear fluorescence for Dorsal and Twist in Dorsal overexpressing embryos as compared to controls (Figure S1B). Further, we investigated mitochondrial dynamics using mito-GFP in live embryos overexpressing Dorsal mounted in end-on orientation (Video 4). 83% (5/6) of the embryos did not form the VF, and the remaining 17% (1/6) of the embryos contained invaginating cells with an abnormal furrow. We compared mitochondrial dynamics in the early time points of gastrulation. We observed that the apical increase in mitochondrial fluorescence in controls was restricted to approximately 8 cells. In Dorsal overexpressing embryos, 33% (2/6) of the embryos showed increased apical mitochondrial fluorescence throughout the embryo, 50% (3/6) showed enhanced apical fluorescence in approximately half of the embryo, and 17% (1/6) of the embryos showed a restricted increase in apical mitochondrial signal (Figure 3D and S1C). We quantified apical:basal mitochondrial intensity in gastrulation when apical constriction was visible and plotted this with respect to the distance along the dorso-ventral axis (Figure 3F). Control embryos showed a peak of apical mitochondrial fluorescence in the ventral region, whereas the mitochondrial fluorescence in Dorsal overexpressing embryos was significantly spread across a greater number of cells. These data showed that Dorsal regulates apical mitochondrial migration in the VF cells.

### Drp1 mutant embryos show accumulation of larger mitochondria in basal regions and decreased apical mitochondrial migration

Loss of mitochondrial fission leads to an increase in mitochondrial size, which affects their migration within cells. Loss of mitochondrial fission protein, Drp1, in neurons has been shown to cause mitochondrial accumulation in the cell body and depletion of mitochondria from synapses and dendrites (Li et al., 2004; Rikhy et al., 2007; Verstreken et al., 2005). Drp1 depleted migrating cells show perinuclear accumulation of mitochondria and defects in actin cytoskeletal remodelling at the leading edge (Desai et al., 2013; Wang et al., 2015; Zhao et al., 2013). Drp1 null mutant germ line clones do not produce embryos(Mitra et al., 2012). We have previously expressed GTPase domain mutants of Drp1 to abrogate mitochondrial fission in embryonic development and neural stem cells (Chowdhary et al., 2020; Dubal et al., 2022). We expressed the GTPase domain mutant of Drp1, Drp1^SG^, to disrupt mitochondrial fission in embryos using *nanos*-Gal4, UASp-mito-GFP. We found that approximately 84% of Drp1^SG^ embryos did not hatch into larvae at 24 hrs. Mitochondria were present in large, clustered spots in the basal regions and small, dispersed spots in apical regions in Drp1^SG^ embryos (78%) compared to small, dispersed mitochondria in apical and basal regions in controls in VF cells (Figure 4A, white arrowhead). We assessed the localization and expression levels of Drp1 in the control and Drp1^SG^ embryos (Figure 4B-D and S2A). In control embryos, Drp1 antibody fluorescence was present on mitochondria in cellularization (Figure 4B, white arrowhead). In Drp1^SG^ expressing embryos, Drp1 antibody fluorescence was present as punctate structures on the clustered mitochondria in the basal sections, and Drp1 punctae did not colocalize with mitochondria in apical regions (Figure 4B, white arrowheads). Drp1 levels were found to be higher in western blots in gastrulation, but this was not statistically significant (Figure 4C). Drp1^SG^ embryos showed a significantly higher expression of Drp1 compared to controls (Figure 4D).

**Figure 4.**
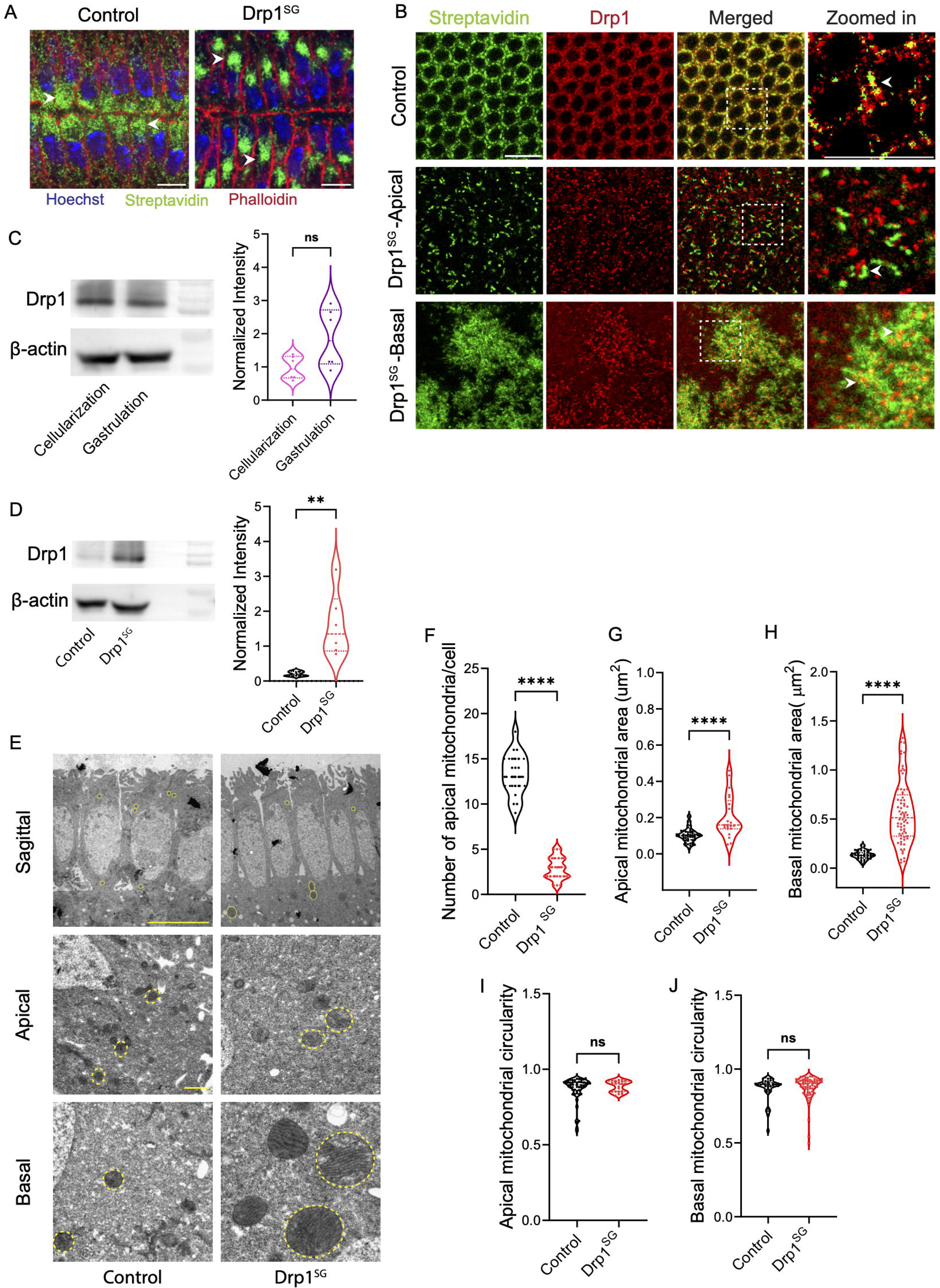
Drp1^SG^ embryos show basal clustering and reduced apical translocation of mitochondria. (A) Representative images of control and Drp1^SG^ embryos with VF, stained for mitochondria (fluorescent streptavidin, green), F-actin (fluorescent phalloidin, red) and DNA (Hoechst, blue) are shown. The white arrowhead marks the mitochondria in the respective embryos. (B-D) Drp1 localization and expression in control and Drp1^SG^ embryos. (B) Control and Drp1^SG^ cellularizing embryos are stained with Drp1 antibody (red) and for mitochondria (fluorescent streptavidin, green). (C) The western blot and graph show Drp1 in control cellularization and gastrulation embryos (N=3). β-actin is used as the loading control. (D) The western blot and graph show Drp1 in control and Drp1^SG^ embryos, with β-actin used as a loading control (N=3). (E) Representative TEM images of late cellularizing control and Drp1^SG^ embryos are shown in sagittal view (1000x magnification) and cross-sectional view (5000x magnification). Mitochondria are electron-dense structures, highlighted with yellow ROI. (F) The plot shows a quantification of mitochondria above the nucleus per cell in control (n=4) and Drp1^SG^ (n=4)embryos. (G) The plot shows the size of mitochondria in apical regions in control (52 mitochondria from 4 embryos) and Drp1^SG^ embryos (21 mitochondria from 4 embryos). (H) The plot shows the size of mitochondria in basal regions in control (34 mitochondria from 4 embryos) and Drp1^SG^ embryos (68 mitochondria from 4 embryos). (I) The plot shows the circularity of mitochondria in apical regions in control (52 mitochondria from 4 embryos) and Drp1^SG^ embryos (21 mitochondria from 4 embryos). (J) The plot shows the circularity of mitochondria in basal regions in control (34 mitochondria from 4 embryos) and Drp1^SG^ embryos (68 mitochondria from 4 embryos). The violin plots show median and quartiles. The statistics were performed by two-tailed unpaired Mann-Whitney test, ns p≥0.05, **p<0.01, ****p<0.0001. Scale Bar: 10μm (A,B), 5μm (E, sagittal), 1μm (E, apical, basal).

We performed transmission electron microscopy of Drp1^SG^ embryos to assess the change in mitochondrial morphology. The mitochondria in late cellularizing Drp1^SG^ embryos were larger compared to controls in apical and basal regions, and basal mitochondria were larger than apical mitochondria (Figures 4E and 4G-4H). Fewer mitochondria were found in apical regions in Drp1^SG^ embryos compared to controls (Figures 4E and F), consistent with immunostainings (Figure 4B). The circularity of mitochondria was similar in Drp1^SG^ expressing embryos and controls (Figures 4I and J). These data showed that mitochondrial fission regulated by Drp1 was essential for regulating mitochondrial size and transport to apical regions in VF cells during gastrulation.

### Drp1 mutant embryos show decreased apical constriction and Myosin II in the ventral cells leading to a broader ventral furrow

VF formation and invagination are facilitated by the apical constriction of ventral cells that follows a ratchet-like mechanism, with repeated pulses of constriction and stabilization resulting in an incremental reduction in the apical cell area. These contractions are driven by an intact acto-myosin network active apically in the VF cells. Starting with discrete spots that intensify in the apical regions across VF cells, and pull the adherence junctions together, Myosin II pulses help to constrict the VF cells (Martin et al., 2010, 2009). We assessed the extent of apical constriction and Myosin II distribution in the ventral cells in Drp1^SG^ embryos expressing reporters of Myosin II light chain, Sqh-GFP and plasma membrane, mScarletCAAX. Drp1^SG^ embryos showed less apical constriction in ventral cells compared to controls, with a similar furrow invagination (Figures 5A-C, Videos 5 and 6). We compared the apical area and Myosin II intensity over time in 40 cells from one control and Drp1 mutant embryo. We observed a progressive decrease in apical area and an increase in Myosin II intensity in controls (Figures 5B and D). However, Drp1^SG^ embryos showed an increase in the number of cells with reduced apical constriction and Myosin II levels (Figure 5B and D). The overall apical area and Myosin II intensity were significantly reduced in Drp1^SG^ embryos compared to controls (Figure 5C and E). We then visualized the overall VF morphology by imaging the embryos in end-on orientation. We observed that Drp1^SG^ embryos had a significantly broader VF with a width of 12.4 (±2.33) μm compared to 8.6 (±0.23) μm in controls (Figures 5F and G, Videos 7 and 8). The apical Myosin II intensity of VF cells in the end-on view was also decreased in Drp1^SG^ embryos compared to controls (Figures 5F and H). These data suggested that reduced apical Myosin II in Drp1^SG^ embryos leads to an overall decrease in apical constriction in VF cells, resulting in a broader VF.

**Figure 5.**
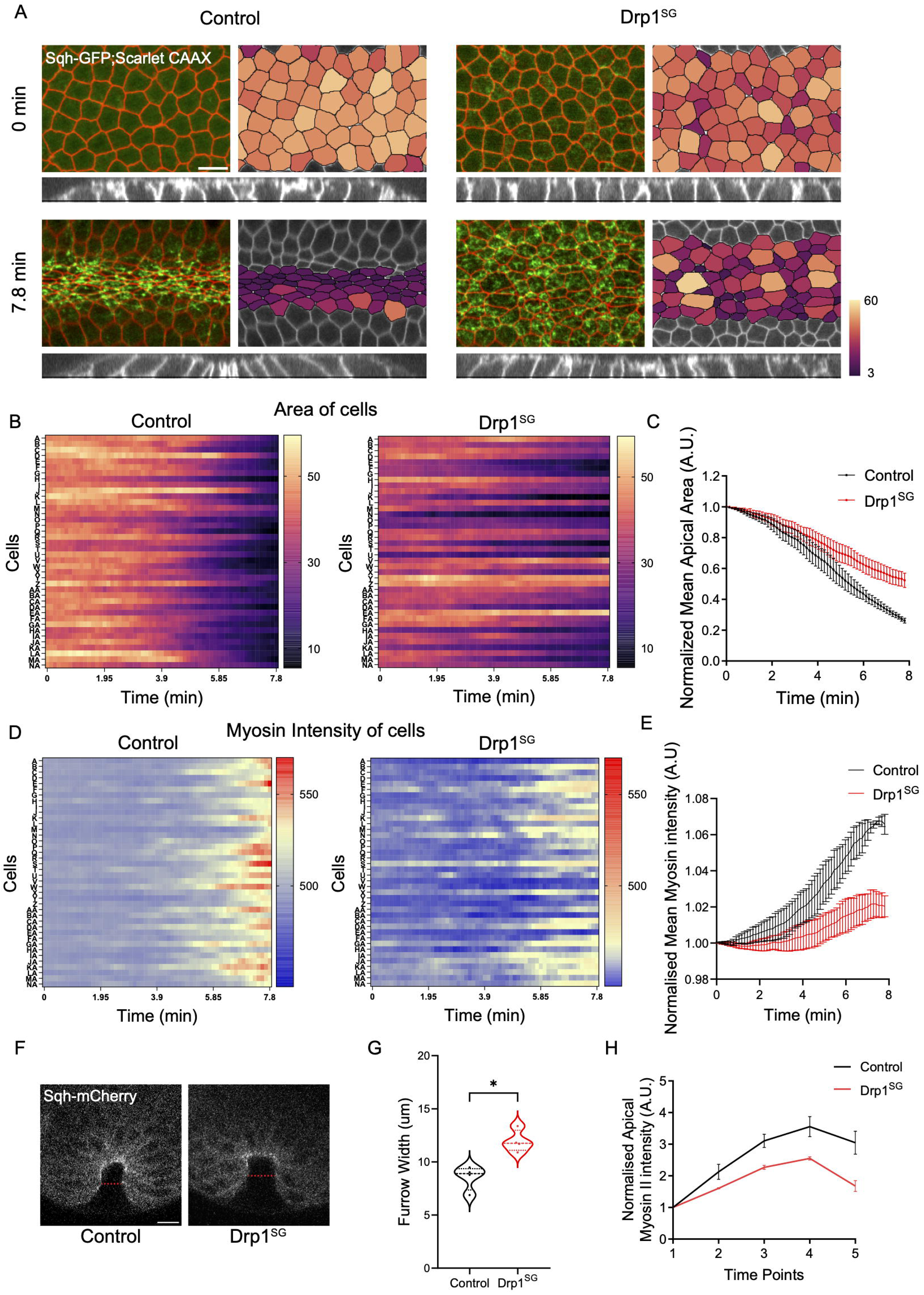
Drp1^SG^ embryos show decreased apical constriction and Myosin II in the ventral cells resulting in a broader ventral furrow. (A) Representative images of live control and Drp1^SG^ embryos, expressing SqhGFP;ScarletCAAX, undergoing apical constriction in ventral cells are shown. Orthogonal views show the extent of furrow ingression. Cells are color coded (right side images) based on area. (B) Heat map of one representative control and Drp1^SG^ embryo shows the area of 40 cells (y axis: A-NA) with time. (C) The plot shows the normalized mean apical area of control and Drp1^SG^ embryos (n=3 embryos each, 40 cells from each embryo). Data is represented as Mean ± SEM. (D) Heat map of one representative control and Drp1^SG^ embryo shows the Myosin II intensity of 40 cells (y axis: A-NA) with time. (E) The plot shows the normalized mean apical Myosin II intensity of control and Drp1^SG^ embryos (n=3 embryos each, 40 cells from each embryo). Data is represented as Mean ± SEM. (F) Representative images, in end-on view, of live control and Drp1^SG^ embryos expressing Sqh-mCherry during VF formation are shown. The red dotted line shows the furrow width measurement. (G) The plot shows the width of the VF in control (n=4) and Drp1^SG^ (n=4). (H) The plot shows the normalized mean myosin II intensity during VF formation in control (n=3) and Drp1^SG^ (n=3). Data is represented as Mean ± SEM. The violin plot shows median and quartiles. The statistics were performed by two-tailed unpaired Mann-Whitney test, *p<0.05. Scale bar 10μm (A,F).

### RNA seq analysis shows no change in transcription of Dorsal/NFkB pathway genes and an increased expression of anti-oxidant genes in Drp1 mutant embryos

Since we found that the Dorsal signaling pathway regulated mitochondrial dynamics in early embryos (Figure 3), we evaluated whether the expression of Dorsal signaling pathway genes and mesodermal genes was affected in Drp1^SG^ expressing embryos, by RNA sequencing. We sorted control and Drp1^SG^ expressing embryos in cellularization and early gastrulation for RNA extraction for sequencing. The principal component analysis from the RNA sequencing output showed clustering of the three replicates of control and Drp1^SG^, and separation between the two genotypes (Figure S3A). In total, 1099 genes were differentially expressed in Drp1^SG^ when compared with control embryos (p value ≤0.05 and Log2FC≤-1 and ≥1). Amongst the differentially expressed genes, 708 genes were up-regulated and 391 genes were down-regulated (Figure 6A). Gene Ontology (GO) analysis performed on these differentially expressed genes was categorized into Biological Process (BP) and Molecular Function (MF). BP enrichment analysis revealed that upregulated genes were associated with ‘biological regulation’, ‘multicellular organismal processes’ and ‘developmental processes’. This is in accordance with another study in mice, where it has been shown that differentially expressed genes (DEGs) in mitochondrial fission depleted embryonic stem cells are enriched in ‘multicellular organism development’ and ‘cell differentiation’ (Seo et al., 2020). Downregulated genes showed enrichment for pathways related to ‘response to stimulus’, ‘locomotion’ and ‘nervous system processes’. MF enrichment analysis showed that upregulated genes were involved in ‘catalytic activity’, ‘ion binding’ and ‘DNA-binding transcription factor activity’. Downregulated genes were enriched in ‘transmembrane transporter activity’, ‘signaling receptor activity’ and ‘molecular transducer activity’ (Figure S3B). We examined the expression of Dorsal signaling pathway genes and found no difference in their expression between control and Drp1^SG^ embryos (Figure 6B). Further using immunostaining, we observed that the nuclear localization of Dorsal and Twist signaling was unaltered in Drp1^SG^ embryos (Figure 6C). We also found that the expression of mesodermal genes, *SPE*, *T48*, *eve* and *MEF2* was unchanged in Drp1^SG^ mutant embryos as compared to controls (Figure 6D). The expression of genes regulating the activation of Myosin II, *Rho1, Rok, RhoGEF2* and *dia* was also similar in both controls and Drp1^SG^ embryos (Figure S3C).

**Figure 6.**
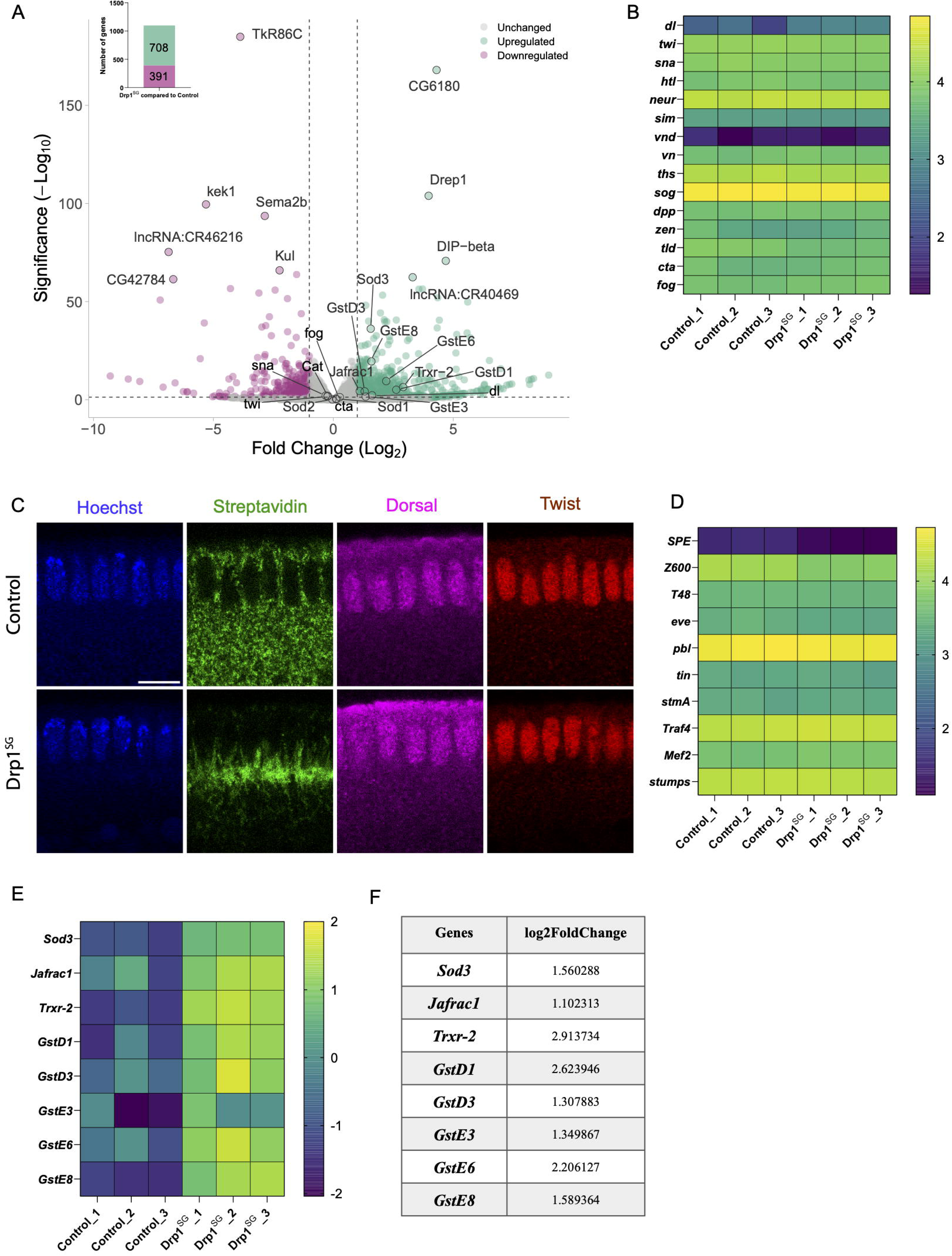
Drp1^SG^ embryos show no change in Dorsal pathway genes, but increased expression of anti-oxidant genes. (A) Volcano plot shows the distribution of genes with the log2foldchange on x axis and significance (-log10pvalue) on y axis. Every point in the plot represents a single gene. All the genes with p value>0.05 and log2foldchange between -1 to +1 are similar in expression in control (N=3) and Drp1^SG^ (N=3), and are marked in grey. Genes marked in green represent the significantly upregulated genes and those in pink indicate the significantly downregulated genes. Top 10 differentially expressed genes (DEGs) and some other genes of interest are highlighted. Bar plot showing the number of significantly upregulated and downregulated genes is represented on the top left corner. (B) Heat map shows the relative expression of Toll-Dorsal pathway genes in control and Drp1^SG^ embryos. (C) Representative sagittal views of control and Drp1^SG^ cellularizing embryos stained for Dorsal (magenta), Twist (red), mitochondria (green) and DNA (blue) are shown. (D) Heat map shows the relative expression of mesodermal genes in control and Drp1^SG^ embryos. (E) Heat map shows the relative expression of anti-oxidant genes in control and Drp1^SG^ embryos. (F) Table shows the log2FoldChange of the anti-oxidant genes in Drp1^SG^ compared to control.

Mitochondria can modulate nuclear gene expression via metabolites such as acetyl-CoA and NAD+, which contribute to epigenetic remodeling. Communication between mitochondria and nucleus is also mediated by ATP, reactive oxygen species (ROS) and Ca^2+^ (Seto and Yoshida, 2014). Importantly, we observed that the expression of anti-oxidant genes such as *Sod3*, *Jafrac1*, *Trxr-2*, *GstD1*, *GstD3*, *GstE3*, *GstE6* and *GstE8* was up-regulated in Drp1^SG^ (Figures 6E and 6F). In addition, using quantitative RT-PCR, we were able to verify an increase in the expression of antioxidant genes *Sod3*, *Jafrac1* and *GstD1* in Drp1^SG^ embryos (Figure S3D).

In summary, the Dorsal signaling pathway was upstream of mitochondrial dynamics in regulating the VF for gastrulation. Change in mitochondrial fission led to an increase in levels of anti-oxidant gene transcription, but no change in Dorsal signaling pathway genes.

### Proteomic analysis shows an increased level of anti-oxidant proteins and a decreased level of ETC complexes in Drp1 mutant embryos

We performed proteome analysis using mass spectrometry of total and mitochondrial protein extracts from control and Drp1^SG^ expressing embryos to assess the change in anti-oxidant proteins and ETC proteins. The principal component analysis showed that the three replicates of control and Drp1^SG^ clustered together, with distinct separation between the two groups (Figure S4A). In the total proteomics, 3838 proteins were detected, out of which 131 were found only in control embryos and 53 were detected only in Drp1^SG^ embryos. 729 proteins were differentially expressed (p value ≤0.05 and Log2FC≤-0.3 and ≥0.3) with 312 being upregulated and 417 being downregulated (Figure 7A). GO analysis was performed on the differentially expressed proteins. It was categorized into Biological Process (BP) and Molecular Function (MF). BP enrichment analysis revealed that upregulated proteins were associated with ‘cellular localization’ and ‘protein transport’. Downregulated proteins showed enrichment for pathways related to ‘metabolic processes’, ‘translation’ and ‘mitochondrial gene expression’. MF enrichment analysis showed that upregulated proteins were involved in ‘catalytic activity’ and various ‘transmembrane transporter activity’. Downregulated proteins were enriched in ‘oxidoreductase activity’ and ‘NADH dehydrogenase activity’ (Figure S4B). We found that anti-oxidant proteins, Sod1, GstD3, GstE1, GstE6, GstE7 and Prx6c were upregulated in Drp1^SG^ (Figures 7B and C).

**Figure 7.**
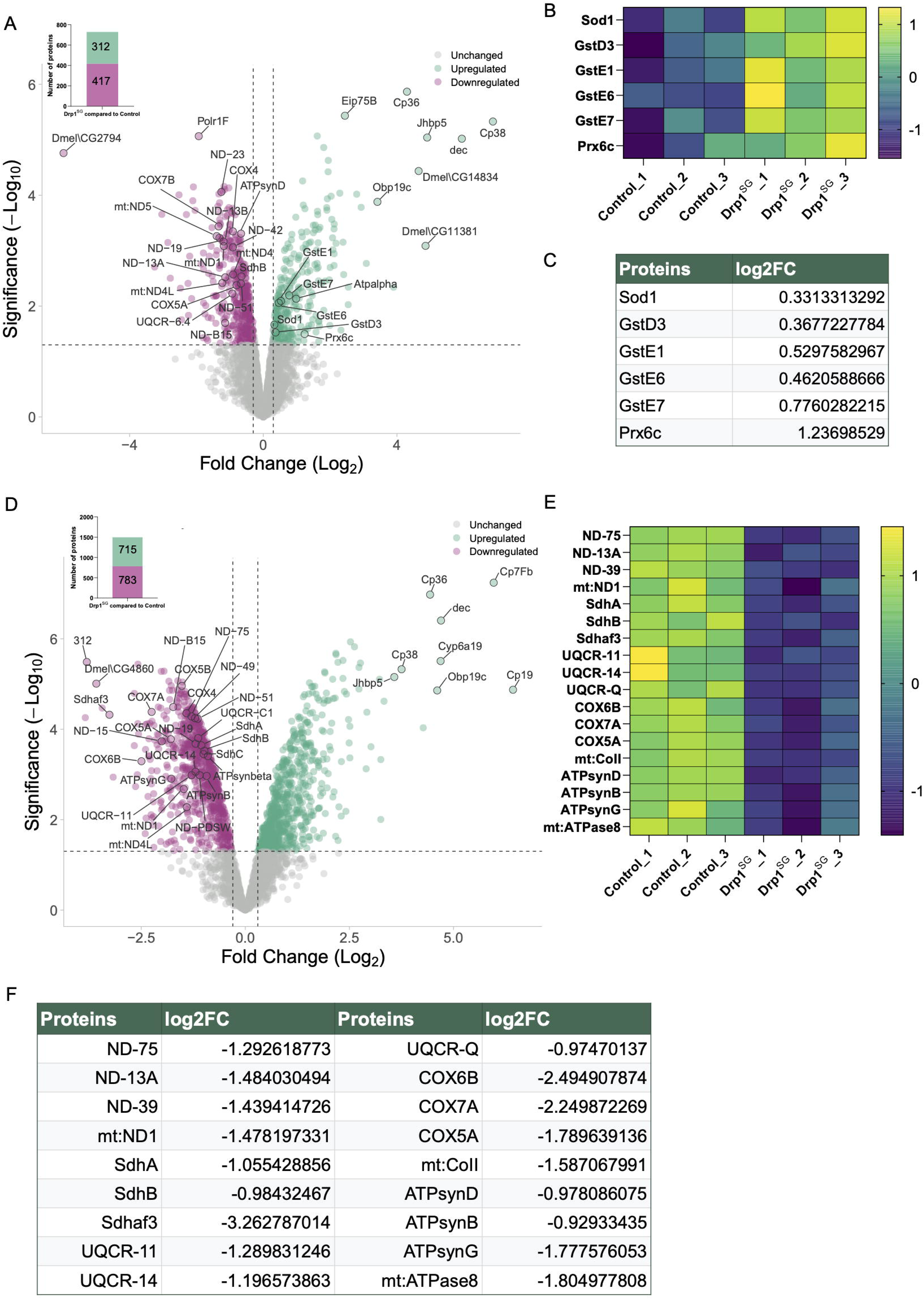
Drp1^SG^ embryos show increased levels of anti-oxidant proteins and downregulation of ETC complexes. (A) Volcano plot shows the distribution of all the proteins identified in the total protein mass spec, with the log2foldchange on the x axis and significance (-log10pvalue) on the y axis. Every point in the plot corresponds to a single protein. All the proteins with p value>0.05 and log2foldchange between -0.3 to +0.3 are marked in grey and are similar in expression in control (N=3) and Drp1^SG^ (N=3) embryos. Green dots represent the upregulated proteins, and pink dots represent the downregulated proteins. Top 10 differentially expressed proteins and some other proteins of interest are highlighted. Bar plot showing the number of significantly upregulated and downregulated proteins is represented on the top left corner. (B) Heat map shows the relative expression of anti-oxidant proteins in control and Drp1^SG^ embryos. (C) Table shows the log2FoldChange of the anti-oxidant proteins in Drp1^SG^ compared to control. (D) Volcano plot shows the distribution of all the proteins identified in the mitochondrial protein mass spec. log2foldchange has been plotted on the x axis and significance (-log10pvalue) has been plotted on the y axis. Every point in the plot represents a protein. All the proteins with p value>0.05 and log2foldchange between -0.3 to +0.3 are similar in expression in control (N=3) and Drp1^SG^ (N=3) and are marked in grey. Upregulated proteins are marked in green and downregulated proteins are marked in pink. Top 10 differentially expressed proteins and some other proteins of interest have been highlighted. Bar plot showing the number of significantly upregulated and downregulated proteins is represented on the top left corner. (E) Heat map shows the relative expression of some of the subunits of all five respiratory complexes in control and Drp1^SG^ embryos. (F) Table shows the log2FoldChange of the subunits of all five respiratory complexes in Drp1^SG^ compared to control.

A total of 3303 proteins were detected from mitochondrial protein extracts, out of which 30 were found only in control embryos and 126 were found only in Drp1^SG^ embryos. Out of the 3147 common proteins, 1498 proteins were differentially expressed with p value ≤0.05 and Log2FC≤-0.3 and ≥0.3. Amongst the differentially expressed proteins, 783 were downregulated and 715 proteins were upregulated (Figure 7D). GO terms were categorized into Biological Process (BP) and Molecular Function (MF) domains. In the BP analysis, amongst the upregulated proteins, there was an enrichment for ‘biological regulation’ and ‘cellular localization and transport’. Downregulated proteins were associated with ‘metabolic processes’ and ‘mitochondrial gene expression, translation and organization’. MF enrichment analysis revealed that upregulated genes were enriched in ‘protein binding’ and ‘catalytic activity’. The downregulated proteins were involved in ‘catalytic activity’, ‘oxidoreductase activity’, and ‘electron transfer activity’ (Figure S4C). Interestingly, we found that many of the subunits of the five ETC complexes were significantly downregulated in the mitochondria of Drp1^SG^ (Figures 7E and 7F), suggesting that the larger mitochondria are metabolically less active. A similar downregulation of ETC complex subunits was observed in the mass spectrometry of total proteins from Drp1^SG^ embryos as well (Figure 7A). Additionally, the levels of ATPbeta in mitochondria isolated from control and Drp1^SG^ expressing embryos using western blot analysis were confirmed to be significantly reduced in Drp1^SG^ (Figure S4D).

Together, the mass spectrometry analysis revealed an increase in anti-oxidant protein levels and a decrease in mitochondrial ETC components, suggesting a decrease in mitochondrial activity in Drp1^SG^ embryos.

### Drp1 mutant embryos show reduced levels of ROS and Drp1^SG^;SOD2^i^ embryos show rescue of anti-oxidant levels, ROS and mitochondrial morphology

ROS is produced in the mitochondria as a result of ETC activity primarily at complex I and complex III (Murphy, 2009). We saw that anti-oxidant gene transcription and protein levels were increased, and ETC subunits from many of the respiratory complexes were reduced in Drp1^SG^ embryos. We therefore estimated the levels of ROS in Drp1^SG^ embryos using the fluorescent dye dihydroethidium (DHE). DHE gets oxidised by superoxides forming a fluorescent product that gives red fluorescence visualized using microscopy (Zhao et al., 2003). We found that the mean fluorescence intensity of DHE was significantly lower in Drp1^SG^ compared to controls (Figures 8A and B). Superoxides formed by the ETC get reduced to H_2_O_2_ with the mitochondrial superoxide dismutase SOD2, and H_2_O_2_ can diffuse out of the mitochondria. Superoxides can also transit into the cytoplasm via the voltage gated anion channel (VDAC) from the mitochondria to the cytoplasm (Han et al., 2003). We checked if there was an increase in overall ROS in embryos depleted of the anti-oxidant mitochondrial superoxide dismutase (SOD2). We found that SOD2 RNAi (SOD2^i^) expressing embryos showed an increase in DHE fluorescence and ROS levels as compared to controls. We made Drp1^SG^;SOD2^i^ combinations and assessed them for ROS and found that the levels were restored and comparable to controls (Figures 8A and B).

**Figure 8.**
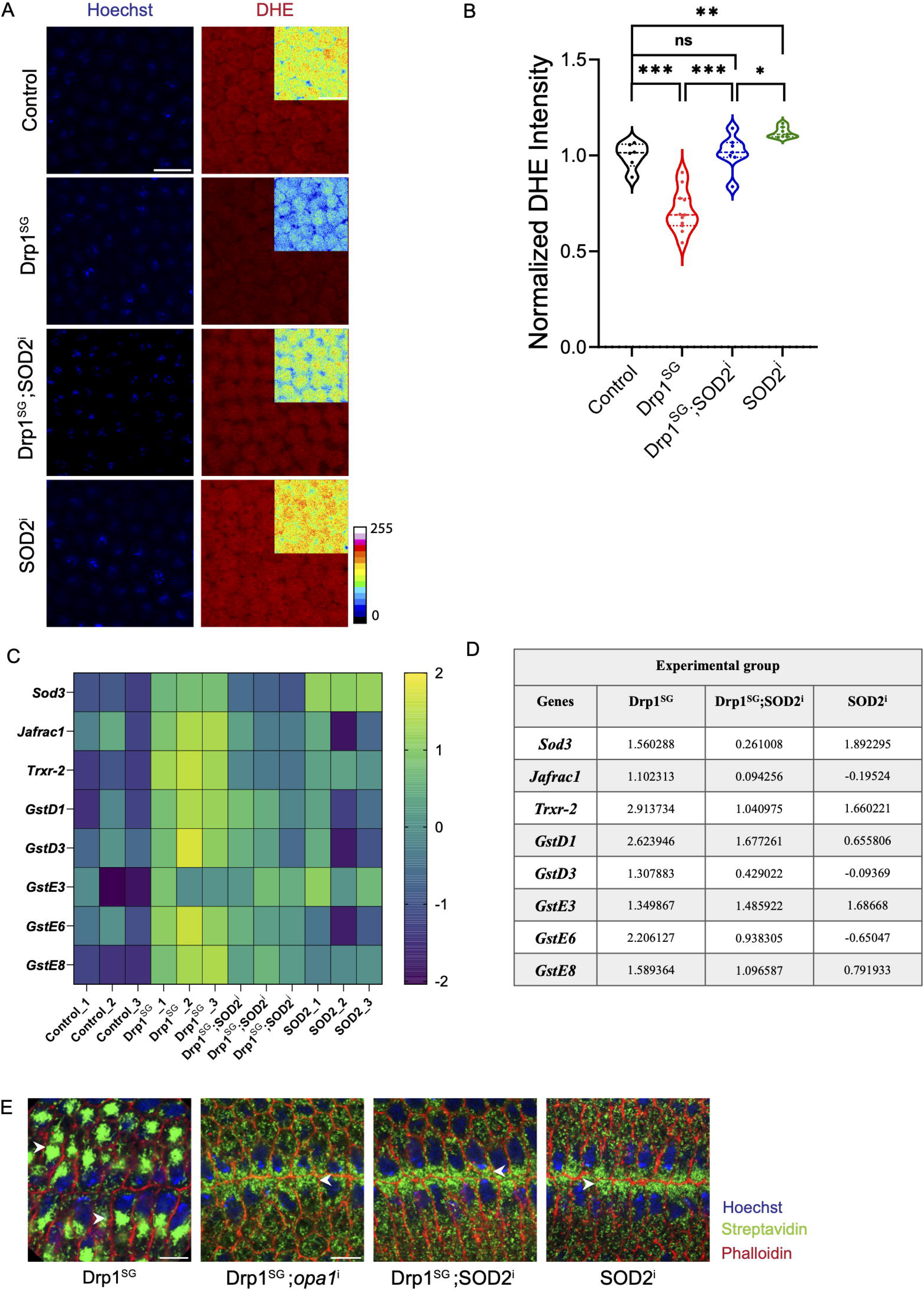
Drp1^SG^ embryos show reduced levels of ROS and Drp1^SG^;SOD2^i^ embryos show rescue of anti-oxidant levels, ROS and mitochondrial morphology. (A) Representative images of control, Drp1^SG^, Drp1^SG^;SOD2^i^ and SOD2^i^ cellularizing embryos, at a single plane just above the nucleus, stained with DHE (red) and Hoechst (blue) are shown. Zoomed in view of the cells is shown in rainbow scale (16 colors) as an inset to highlight the relative difference in intensities. (B) The plot shows the normalized mean fluorescence intensity of DHE. n=6 for control, n=11 for Drp1^SG^, n=7 for SOD2^i^, n=7 for Drp1^SG^;SOD2^i^. (C) Heat map shows the relative expression of anti-oxidant genes in control, Drp1^SG^, Drp1^SG^;SOD2^i^ and SOD2^i^ embryos. (D) Table shows the log2FoldChange values of the anti-oxidant genes, each with respect to control. (E) Representative images of Drp1^SG^, Drp1^SG^;*opa1*^i^ and Drp1^SG^;SOD2^i^ and SOD2^i^ gastrulating embryos with VF stained for DNA (Hoechst, blue), F-actin (fluorescent phalloidin, red) and mitochondria (fluorescent streptavidin, green). White arrowhead marks the mitochondria in respective embryos. The violin plot shows median and quartiles. The statistics were performed by two-tailed unpaired Mann-Whitney test, ns p≥0.05, *p<0.05, **p<0.01, ***p<0.001. Scale Bar 10μm (A,E).

We then asked if combining Drp1^SG^ and SOD2^i^ could alleviate the gene expression changes seen in Drp1^SG^ embryos. Principal component analysis showed that 2 out of 3 samples of Drp1^SG^;SOD2^i^ clustered between control and Drp1^SG^, suggesting a partial reversal of the transcriptional changes (Figure S3A). We examined the expression of anti-oxidant genes that were upregulated in Drp1^SG^ (log2FC > 1 compared to the control). We compared their log2FC values in Drp1^SG^;SOD2^i^ calculated relative to control and found that the genes *Sod3, Jafrac1, Trxr-2, GstD1, GstD3, GstE6* and *GstE8*, but not *GstE3*, showed a reduced log2FC compared to Drp1^SG^ (Figures 8C and D), indicating rescue of their expression levels upon depletion of SOD2 in Drp1 mutant embryos.

ROS has been shown to impact mitochondrial morphology. Increasing ROS leads to mitochondrial fragmentation in *Drosophila* follicle cells and cells undergoing delamination during *Drosophila* dorsal closure (Muliyil and Narasimha, 2014; Uttekar et al., 2024). We therefore checked mitochondrial distribution in Drp1^SG^;SOD2^i^ embryos in the VF. We observed a reduction in basal mitochondrial clustering and a rescue of apical mitochondrial migration in the VF cells in Drp1^SG^;SOD2^i^ embryos as compared to Drp1^SG^ alone (Figure 8E). Oxidative stress causes mitochondrial fragmentation and release of mitochondrial fusion protein, Opa1, into the cytosol in mammalian HT22 cells (Sanderson et al., 2015). Opa1 depletion also causes an increase in ROS in *Drosophila* follicle cells and mitochondrial fragmentation in follicle cells and embryos (Chowdhary et al., 2020; Uttekar et al., 2024). We therefore knocked down Opa1 in Drp1^SG^ embryos and observed that the mitochondrial migration and clustering defect was suppressed (Figure 8E). Together, these data showed that Drp1 mutant embryos have decreased ROS and increasing ROS supported mitochondrial fission for apical transport in VF cells during gastrulation.

### Fragmenting mitochondria and increasing ROS rescues apical Myosin II and ventral furrow formation in Drp1^SG^;*opa1^i^* and Drp1^SG^;SOD2^i^ embryos

Drp1^SG^ embryos showed a decrease in ROS, apical Myosin II and apical constriction (Figures 5 and 8). The Drp1^SG^;SOD2^i^ combination showed increased ROS and rescue of mitochondrial migration to apical regions (Figure 8). ROS has been shown to regulate Myosin II in various processes such as dorsal closure (Muliyil and Narasimha, 2014), wound healing (Hunter et al., 2018) and *Drosophila* cellularization (Chowdhary et al., 2020). We therefore assessed Myosin II levels and VF morphology in Sqh-mCherry expressing Drp1^SG^;*opa1*^i^ and Drp1^SG^;SOD2^i^ embryos (Videos 9 and 10). Drp1^SG^;*opa1*^i^ showed a partial rescue in the apical Myosin II levels, and the furrow width was 6.5 (±0.91) μm, which was restored compared to Drp1^SG^ and even smaller than controls. The apical Myosin II intensity of Drp1^SG^;SOD2^i^ was similar to that of controls, and the furrow width was 7.3 (±1.69) μm, which was significantly smaller compared to Drp1^SG^. SOD2^i^ embryos showed Myosin II levels and furrow width (6.98±0.81 μm) comparable to the controls (Figures 9A-C, Video 11). This data shows that fragmenting mitochondria, either by decreasing fusion or increasing ROS, leads to restoration of apical Myosin II localization and VF formation. In summary, our data collectively show that mitochondria regulate VF furrow dynamics through ROS mediated regulation of Myosin II contractility in VF cells.

**Figure 9.**
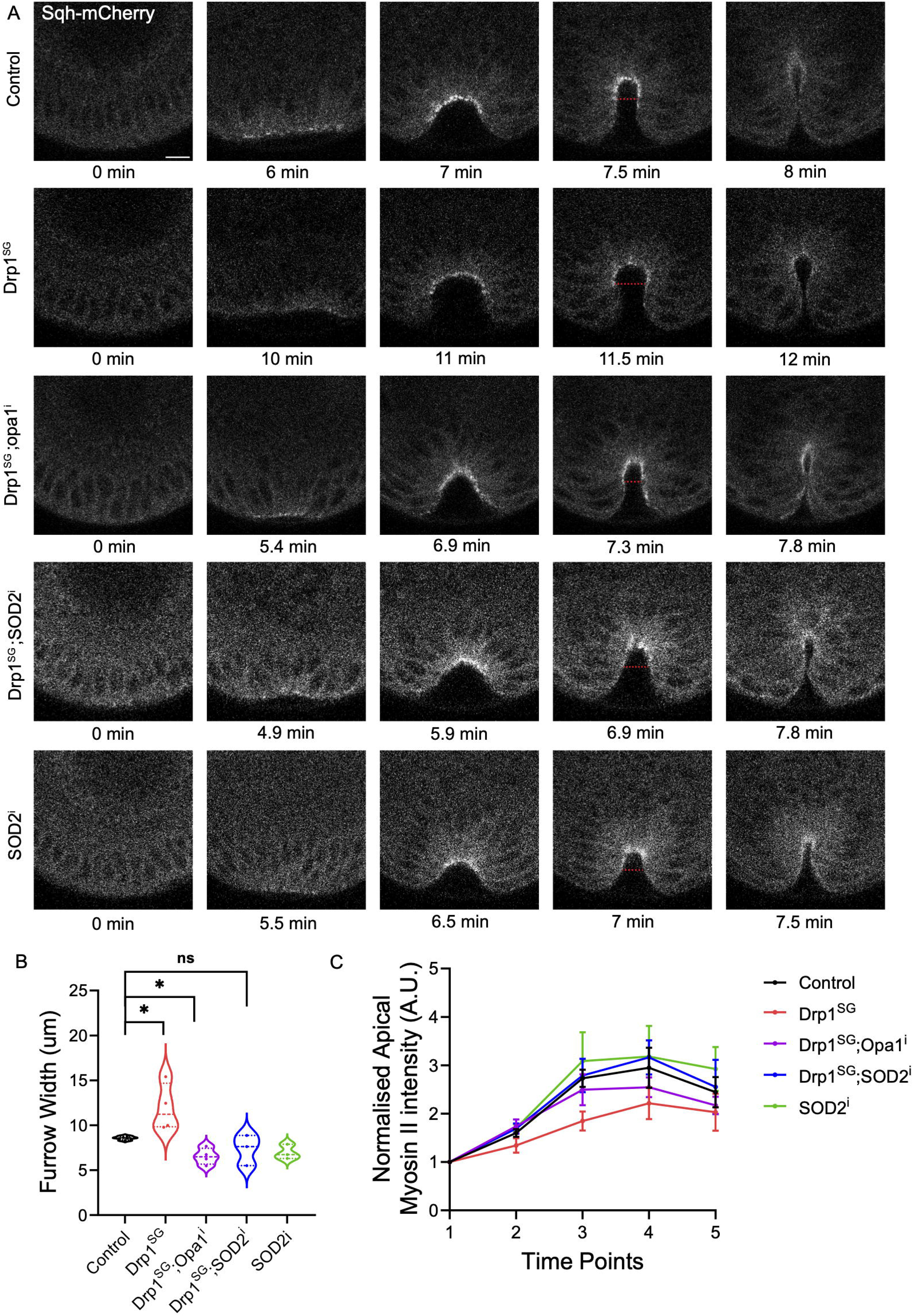
Drp1^SG^;*opa1^i^* and Drp1^SG^;SOD2^i^ embryos show a rescue of apical Myosin II and ventral furrow formation. (A) Representative images in end-on view from live embryos (control, Drp1^SG^, Drp1^SG^;*opa1^i^*, Drp1^SG^;SOD2^i^ and SOD2^i^) expressing Sqh-mCherry, starting from the end of cellularization (0 min) till invagination of the VF are shown. The red dotted line shows the measurement of the furrow width. (B) The plot shows the furrow width, measured at the time point 4 when the VF is U shaped. n=4 for control, n=4 for Drp1^SG^, n=3 for SOD2^i^, n=3 for Drp1^SG^;SOD2^i^, n=4 for Drp1^SG^;*opa1^i^* embryos (C) The plot shows the normalized mean apical fluorescence of Sqh-mCherry in the VF at the 5 time points. n=4 for control, n=4 for Drp1^SG^, n=4 for SOD2^i^, n=3 for Drp1^SG^;SOD2^i^, n=4 for Drp1^SG^;*opa1^i^* embryos. Data is represented as Mean ± SEM. The violin plot shows median and quartiles. The statistics were performed by two-tailed unpaired Mann-Whitney test, ns p≥0.05, *p<0.05. Scale 10μm (A).

**Figure 10.**
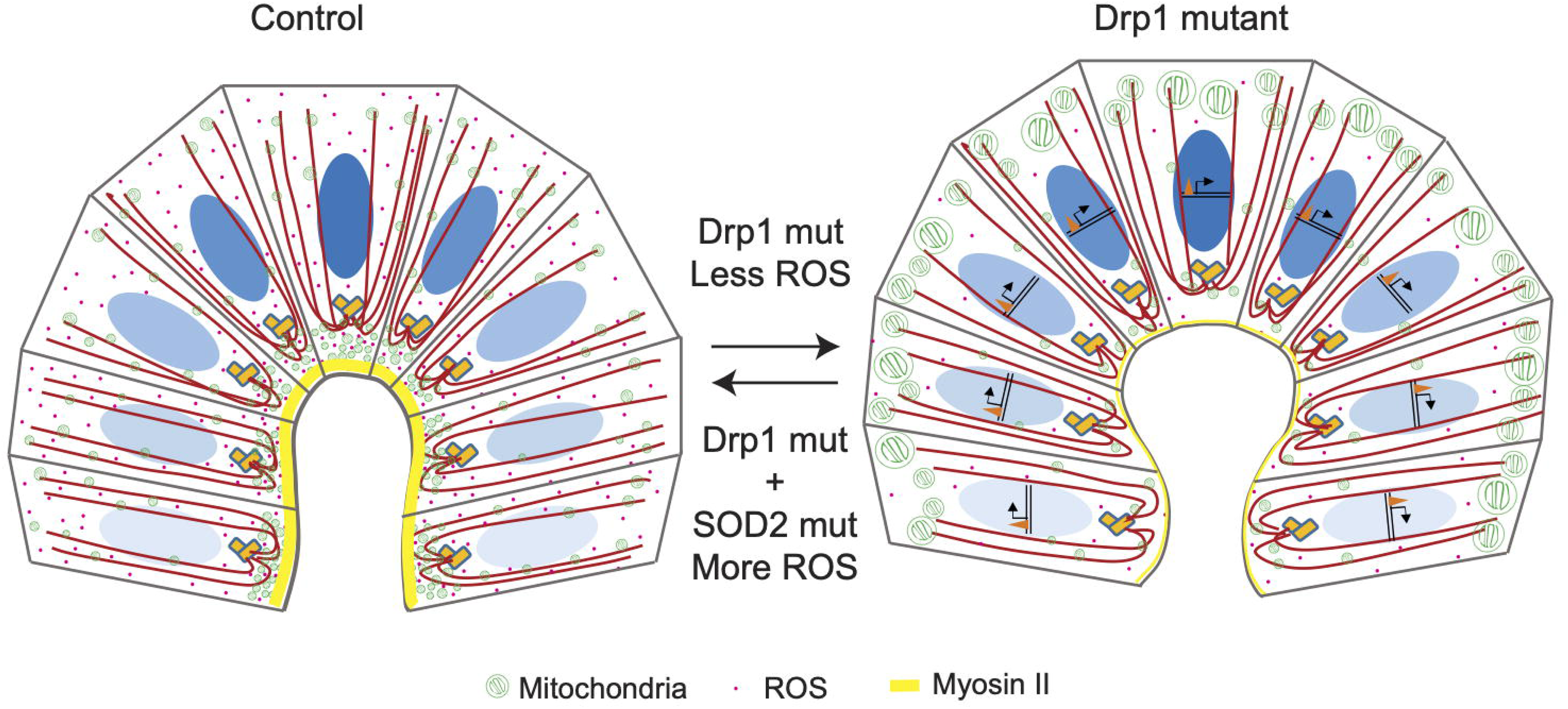
Summary Model. Mitochondria (green) migrate apically in the VF with the help of microtubules (red lines), resulting in their apical enrichment. The apical mitochondrial migration is regulated by Dorsal (blue nuclear gradient). Drp1 mutant embryos (Drp1^SG^) display larger mitochondria, which remain at the basal side (green) and are unable to migrate apically, resulting in few smaller mitochondria in the apical regions. Expression of Dorsal signaling pathway genes and mesodermal genes is not affected in Drp1^SG^. Anti-oxidant genes are upregulated in Drp1^SG^ (shown by transcription of red triangle inside the nucleus) in Drp1^SG^. Levels of ETC complexes subunits and ROS (ROS, pink dots) are reduced in Drp1^SG^. Myosin II levels at the VF are decreased (yellow), resulting in a broader VF. Knockdown of SOD2 and *opa1* in Drp1^SG^ results in fragmentation of mitochondria and rescue of apical mitochondrial migration in the VF. Levels of anti-oxidants and ROS are also rescued in Drp1^SG^;SOD2^i^. Myosin II levels at the VF were restored in both Drp1^SG^;SOD2^i^ and Drp1^SG^;*opa1^i^*, resulting in rescue of the VF formation defect.

## Discussion

We demonstrate that small, punctate and fragmented mitochondria migrate from basal to apical regions in ventral cells during ventral furrow formation in gastrulation in *Drosophila* embryogenesis. Mitochondria migrate to apical regions in all cells during cellularization (Chowdhary et al., 2020). However, migration in gastrulation occurs specifically at the VF in a microtubule-dependent manner and, notably, is regulated by the Dorsal signaling pathway. We find that maintaining mitochondrial size is essential to migrate, as larger mitochondria in Drp1 mutant embryos are unable to migrate apically. Also interestingly, reverting mitochondrial size with SOD2^i^ and *opa1*^i^ in Drp1^SG^ embryos enables them to migrate to apical regions. RNA seq analysis reveals that Drp1 mutant embryos do not show a change in the Dorsal signaling pathway and mesodermal genes, but show an increased expression of anti-oxidant genes. This confirms that the Dorsal signaling pathway is upstream of mitochondrial migration. The mitochondria in Drp1 mutant embryos have reduced activity due to a decrease in the levels of ETC components, leading to a decrease in ROS. Since the embryos also have an increase in anti-oxidant gene transcription, we conclude that they have an overall reduced environment. The apical Myosin II levels are decreased in Drp1 mutant embryos during gastrulation, resulting in reduced apical constriction, leading to larger ventral cells and broader VF. Suppression of these defects upon fragmenting mitochondria and increasing ROS, coincident with mitochondrial migration in the VF cells, indicates that apical mitochondrial migration is essential for this morphogenetic process.

Mitochondria have been shown to exert their effect on the patterning of embryonic axes in various organisms by their differential distribution and local changes in activity. In sea urchin embryos, mitochondrial density and cytochrome oxidase activity at the oral pole are higher, thereby establishing a respiratory gradient along the oral-aboral (OA) axis, which is thought to activate the Nodal signaling pathway (Coffman, 2009; Czihak, 1963). The polarity of the OA axis can be altered by changing the respiratory gradient (Pease, 1942, 1941; Pilocarpine and Cysteine, n.d.), and dorsal activation of HIFα specifically contributes to dorso-ventral (DV) axis specification (Chang et al., 2017). A gradient of mitochondrial membrane potential with higher respiratory activity on the ventral side has also been found in *Drosophila* blastoderm embryos (Schiffmann, 1997). In *Xenopus* embryos, the cells of the dorsal blastopore lip in the Spemann-Mangold organizer region undergo morphological changes in the form of apical constriction and mesoderm internalization similar to the *Drosophila* VF. Multiple studies have reported the dorsal region of the *Xenopus* embryos to have higher mitochondrial content and oxygen consumption (Garfinkel et al., 2023; Landström and Lovtrup, 1975; Løvtrup and Pigon, 1958; Yost et al., 1995). The organizer has been shown to be established by oxidative metabolism driven by mitochondrial activity (Garfinkel et al., 2023). In mouse and zebrafish embryos, mitochondrial ribosomal proteins and mitochondrial ATP synthesis have also been shown to be critical for gastrulation, respectively (Cheong et al., 2020; Pinho et al., 2013). Thus, the common theme that appears from these studies and our findings is that there is a preferential mitochondrial localization and/or activity in the embryonic regions of high cellular dynamics, which is essential for driving cellular remodeling in these regions. Drp1 mutant embryos in our study have reduced activity and metabolites, including ROS. It is also possible that other metabolites, such as NAD+, are decreased, which could not only change the local metabolism but also give rise to epigenetic changes in the genome, further preserving the reduced environment.

An interaction between mitochondria and Fog, a downstream effector in the Dorsal signalling pathway, has been reported previously. Overexpression of Fog, in *Drosophila* third instar larval muscles, leads to mitochondrial fission in a Drp1 dependent manner (Ratnaparkhi, 2013). Fog is also enriched apically in the VF cells (Dawes-Hoang et al., 2007). Thus, Fog activity could maintain mitochondrial fission in ventral cells, facilitating apical mitochondrial migration in the furrow. Downregulation of Drp1 inhibits Fog signalling and also leads to VF defects, similar to Fog loss of function mutants (Ratnaparkhi, 2013). Additionally, different isoforms of Drp1 were reported in ventralized and lateralized embryos during gastrulation in proteomic analysis. Knockdown of Drp1 using dsRNA resulted in mild to severe defects in the VF (Gong et al., 2004). However, the expression of Toll-Dorsal pathway genes and mesodermal genes was unchanged in Drp1^SG^, despite showing VF defects. This indicates that VF invagination and mesoderm formation are regulated independently. It has also been reported in a previous study that VF formation and mesoderm differentiation need not be associated with each other. In the absence of Twist activity, Snail is sufficient to induce VF invagination but not mesoderm differentiation (Ip et al., 1994). It has also been shown in sea urchin embryos that the gut and myocyte progenitors can differentiate even in the absence of invagination in exogastrulae (Ransick and Davidson, 1993; Venuti et al., 1993).

Mitochondrial shape and activity are closely associated with each other. Changes in mitochondrial morphology have been shown to alter ROS levels (Ježek et al., 2018) and affect calcium homeostasis (Kowaltowski et al., 2019). Increased fission can alter cell physiology and promote cancer cell survival by increasing ROS production (Huang et al., 2016). Mitochondrial fission, upon loss of Opa1, leads to an increase in ROS levels (Rubén et al., 2021; Tang et al., 2009; Uttekar et al., 2024; Yarosh et al., 2008). It is conventionally believed that increased mitochondrial fusion supports enhanced ETC activity (Chen et al., 2023; Yao et al., 2019). However, some studies have reported a decrease in ROS and mitochondrial activity upon Drp1 depletion, similar to our finding (Parone et al., 2008; Ponte et al., 2020). It is likely that the bigger mitochondria inherited from oogenesis in Drp1^SG^ embryos could be functionally compromised, resulting in calcium imbalance and decreased ROS production from ETC complexes. ROS can act as a signaling molecule and modify enzymes by oxidising cysteine residues (Moldogazieva et al., 2018), leading to carbonylation of proteins (Fedorova et al., 2009). Calcium can also regulate the cytoskeletal network directly or via regulatory proteins. Myosin II light chain kinase (MLCK) is a serine-threonine kinase that is activated by calmodulin, upon an increase in calcium levels, and further it phosphorylates the regulatory Myosin II light chains, resulting in increased contractility (Robison and Colbran, 2013; Scholey et al., 1980). Intracellular calcium can drive remodeling of F-actin, thereby inducing apical constriction leading to the formation of the neural tube in *Xenopus (Suzuki et al., 2017)*. ROS is known to regulate kinases and phosphatases through oxidation (Corcoran and Cotter, 2013; Ray et al., 2012). ROS has been shown to regulate two key Myosin II regulators, Rho kinase (ROCK) and Src kinase (Hunter et al., 2018; Muliyil and Narasimha, 2014) in *Drosophila* dorsal closure and zebrafish wound healing, respectively. It is possible that the ROS levels in Drp1^SG^;*opa1^i^* might be increased due to mitochondrial fission and thus the increased ROS, in Drp1^SG^;SOD2^i^ and Drp1^SG^;*opa1^i^*, could rescue the Myosin II activity via upstream kinases and/or phosphatases. Thus, we conclude that mitochondria migrate to the apical side in the VF cells to assist in Myosin II activation via ROS in a spatiotemporal manner, thereby aiding VF formation. Additionally, mitochondria could also regulate actin via ROS and/or calcium and could also be essential for providing ATP or other metabolites for driving cell shape changes.

## Acknowledgements

We thank IISER Pune microscopy and *Drosophila* facilities for help with imaging and *Drosophila* stocks and crosses. We thank RR lab members for constant input on this work. We thank FlyBase and Bloomington Stock Centers for the information and stocks used in this study. SM and SC thank the Council of Scientific and Industrial Research CSIR, India and the University Grants Commission UGC for the graduate student fellowship. HM thanks IISER Pune for the graduate student fellowship. We thank J-P. Chauvin, A. Aouane, and F. Richard at the Institut de Biologie du Développement de Marseille (IBDM) electron microscopy facility (Marseille, France) for help with embryo processing and image acquisition. We thank the Transmission Electron Microscope facility, ACTREC, Tata Memorial Centre, Kharghar, Navi Mumbai, India for help with sample processing, sectioning and imaging control and Drp1 mutant embryos. We thank Eric F. Wieschaus (Princeton University, USA), Yu-Chiun Wang (Riken, Kobe, Japan) and Girish Ratnaparkhi (IISER, Pune, India) for antibodies and fly stocks used in this study. We thank V Proteomics, Delhi, India for mass spectrometry analysis and Medgenome, Bangalore, India for RNA seq analysis.

## Author contribution

SM, SC and RR designed the study, performed the experiments and analysed the data. SP and SM performed and analysed the western blots, RTPCR, genomics and proteomics. MM performed the experiment in Figure 1F-G, SC analysed the experiment. HM performed and analyzed the experiments in 4E-J. RR acquired the funding. SM and RR wrote the manuscript. SM, SC, HM, MM and RR edited the manuscript.

## Supplementary Figure Legends

**Figure. S1 Dorsal overexpression increases Dorsal and Twist expression**

(A) Representative images of control and Dl-OE embryos in cellularization, stained for Dorsal, Twist and Hoechst (DNA, blue) are shown. Scale bar 10μm.

(B) The plot shows the Ventral:Dorsal intensity ratio of Dorsal and Twist in control (n=7, 6 respectively) and Dl-OE (n=4, 5 respectively). The violin plot shows median and quartiles. The statistics were performed by two-tailed unpaired Mann-Whitney test, **p<0.01.

(C) Representative snapshots from live embryos overexpressing Dorsal (Dl-OE) and mito-GFP are shown in gastrulation. The 16 color rainbow scale is used to represent fluorescence intensities of mito-GFP on a 8 bit scale where 0 is black and 255 is white.

**Figure S2. Western blot with anti-Drp1**

(A) Western blot with anti-Drp1 in *Drosophila* embryos in cellularization (n=3)

**Figure S3. RNA seq analysis and qPCR of Drp1 mutant embryos**

(A) Principal component analysis (PCA) of control, Drp1^SG^, SOD2^i^, Drp1^SG^;SOD2^i^ embryos in RNA sequencing is shown (n=3 for each genotype).

(B) Gene ontology (GO) analysis of upregulated and downregulated genes in control and Drp1^SG^ is shown.

(C) Heat map shows the relative expression of the genes that regulate Myosin II in control and Drp1^SG^ embryos.

(D) The plot shows the relative mRNA expression of 3 anti-oxidant genes- *Jafrac1*, *Sod3* and *GstD1* (n=3). The statistics were performed by two-tailed unpaired Mann-Whitney test, **p<0.01. ***p<0.001.

**Figure S4. Total proteomics and mitochondrial proteomics analysis of Drp1 mutant embryos**

(A) Principal component analysis of control (C) and Drp1^SG^ (D) embryos in total proteomics and mitochondrial proteomics is shown (n=3 for each genotype).

(B) Gene ontology (GO) analysis of upregulated and downregulated proteins in total proteomics in control and Drp1^SG^ is shown.

(C) Gene ontology (GO) analysis of upregulated and downregulated proteins in mitochondrial proteomics in control and Drp1^SG^ is shown.

(D) Relative expression of ATPbeta in mitochondria extracts of control and Drp1^SG^ embryos is shown using western blots (left). Coomassie staining is shown for equal protein loading (centre). The plot shows the normalized intensity of ATPbeta in control and Drp1^SG^ embryos (n=3). The violin plot shows median and quartiles. The statistics were performed by two-tailed unpaired Mann-Whitney test, ns p≥0.05.

## Video legends

**Video 1.** Mitochondrial dynamics in mito-GFP expressing embryo undergoing ventral furrow formation, shown in 16 color rainbow scale. Time is shown in minutes and seconds, and the scale bar is 10 μm.

**Video 2.** Mitochondrial dynamics and Myosin II distribution in mito-PAGFP, Sqh-mCherry expressing embryo. Time is shown in minutes and seconds, and the scale bar is 10 μm.

**Video 3.** Gastrulation is shown in heterozygous *dl*^1^ mutant embryos expressing mito-GFP, shown in 16 color rainbow scale. Time is shown in minutes and seconds, and the scale bar is 10 μm.

**Video 4.** Gastrulation is shown in UASp-Dl embryos expressing mito-GFP, shown in 16 color rainbow scale. Time is shown in minutes and seconds, and the scale bar is 10 μm.

**Video 5.** Apical constriction at the VF in Sqh-GFP;mScarletCAAX expressing embryos. Time is shown in minutes and seconds, and the scale bar is 10 μm.

**Video 6.** Apical constriction at the VF in Drp1^SG^ embryos expressing Sqh-GFP;mScarletCAAX. Time is shown in minutes and seconds, and the scale bar is 10 μm.

**Video 7.** VF formation is shown in end-on view in Sqh-mCherry expressing embryos. Time is shown in minutes and seconds, and the scale bar is 10 μm.

**Video 8.** VF formation is shown in end-on view in Drp1^SG^ embryos expressing Sqh-mCherry. Time is shown in minutes and seconds, and the scale bar is 10 μm.

**Video 9.** VF formation is shown in end-on view in Drp1^SG^;*opa1*^i^ embryos expressing Sqh-mCherry. Time is shown in minutes and seconds, and the scale bar is 10 μm.

**Video 10.** VF formation is shown in end-on view in Drp1^SG^;*SOD2*^i^ embryos expressing Sqh-mCherry. Time is shown in minutes and seconds, and the scale bar is 10 μm.

**Video 11.** VF formation is shown in end-on view in *SOD2*^i^ embryos expressing Sqh-mCherry. Time is shown in minutes and seconds, and the scale bar is 10 μm.

