## Supplementary figures and images for "Spatiotemporal localization and activity of mitochondria regulate actomyosin contractility in *Drosophila* gastrulation"

### Figure S1, Figure S2, Figure S3, Figure S4

Supplementary Fig S1

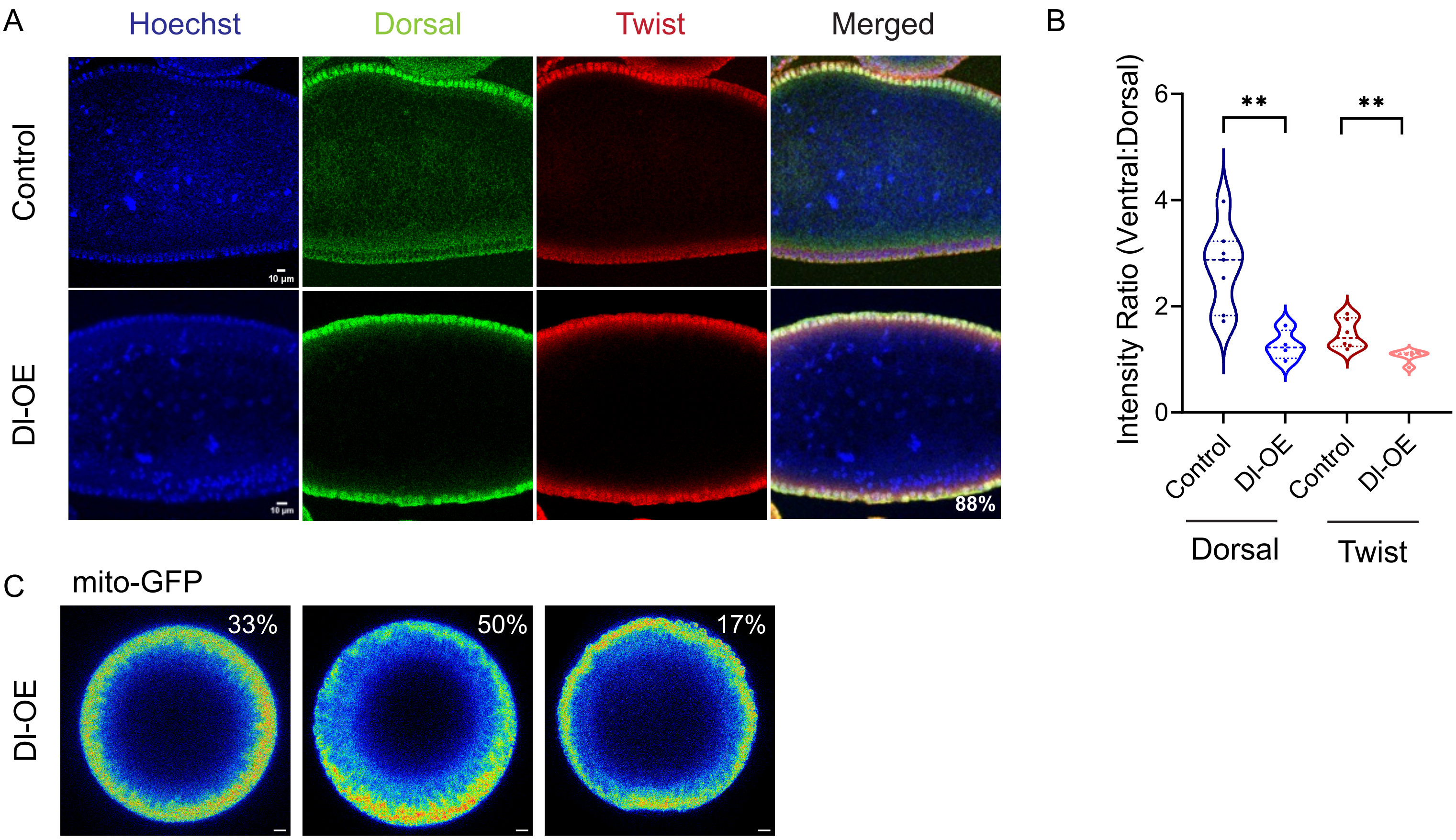

Supplementary Fig S2

A

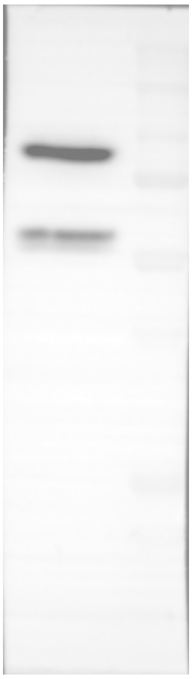

Supplementary Fig S3

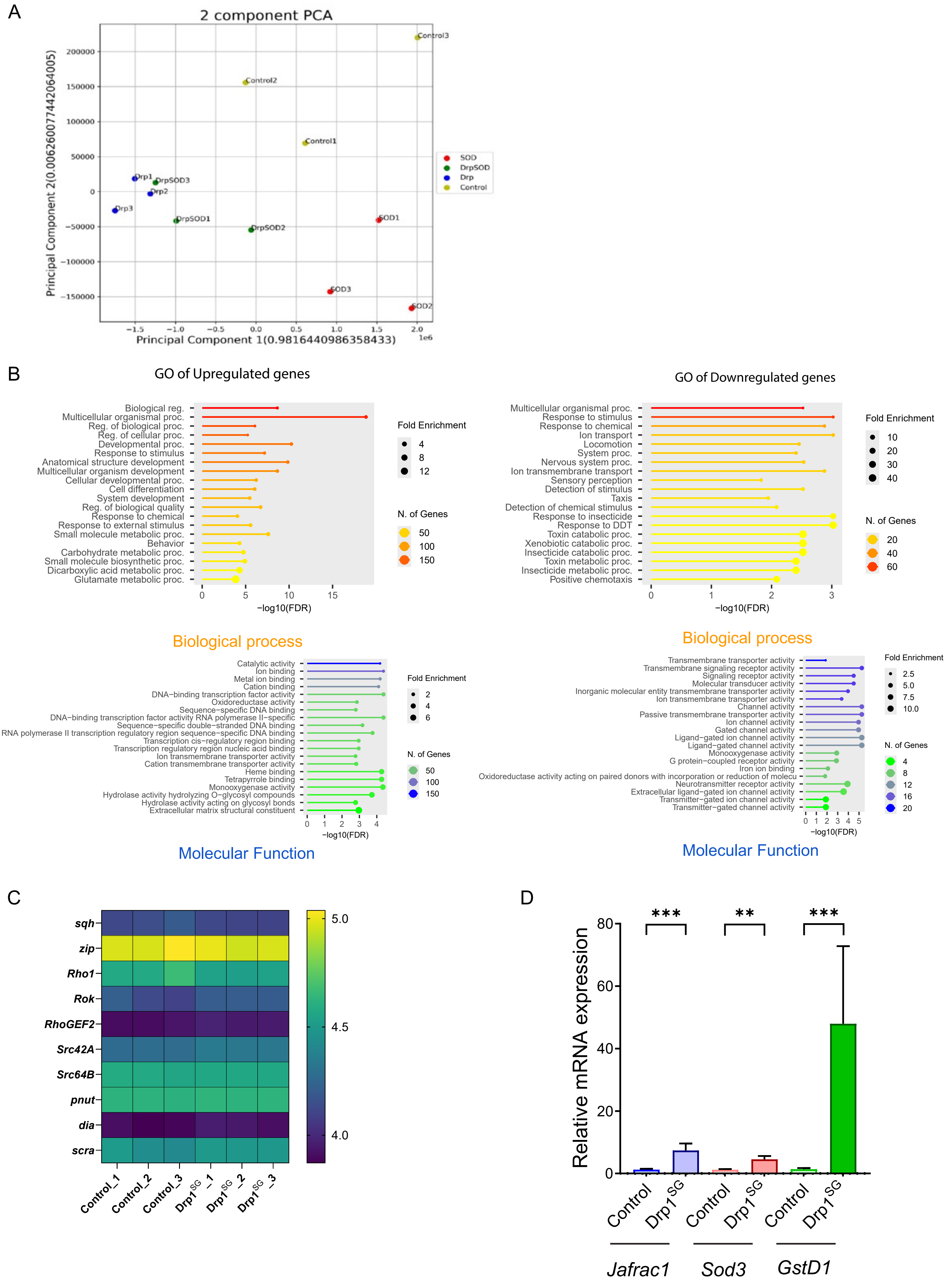

Supplementary Fig S4

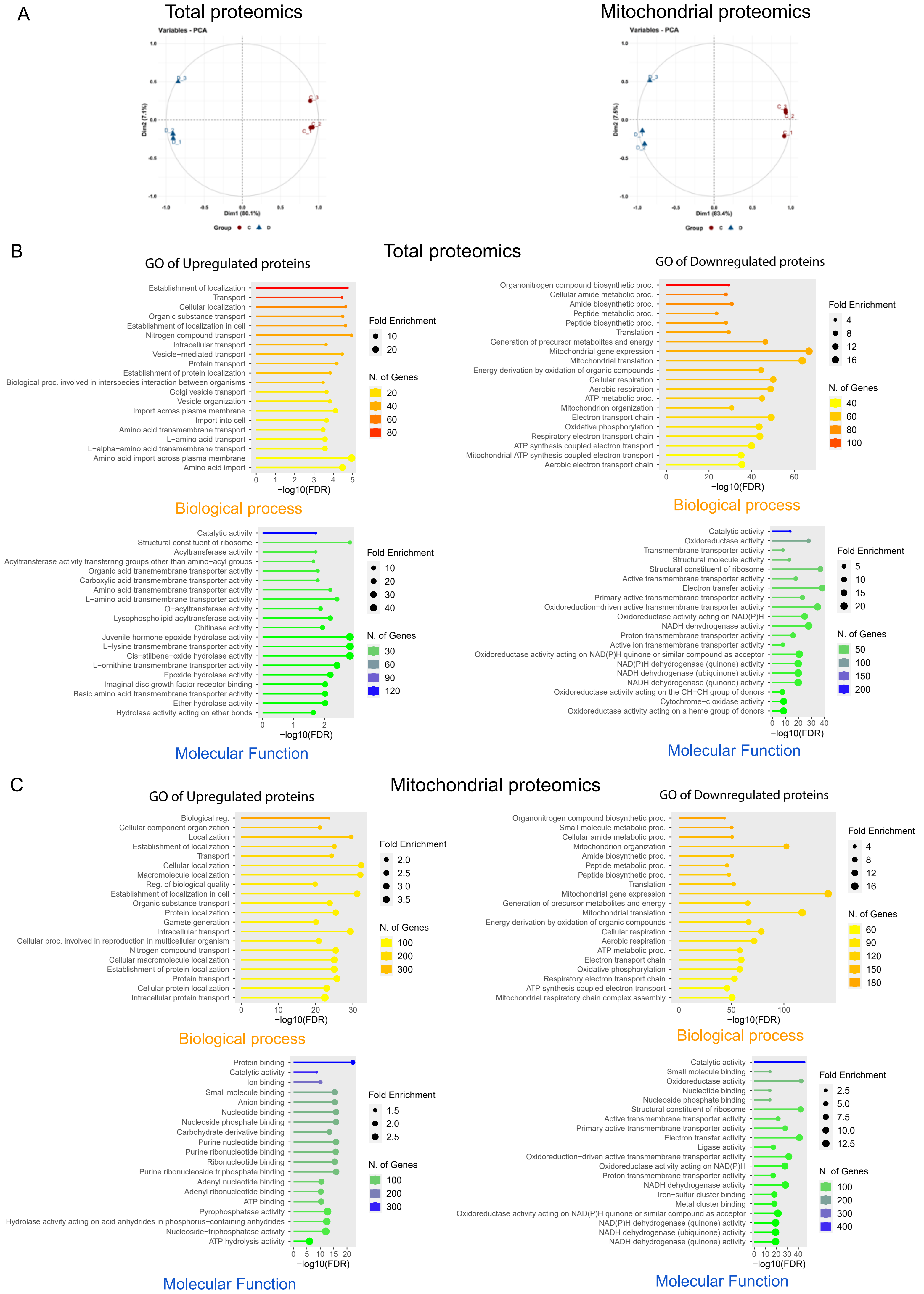

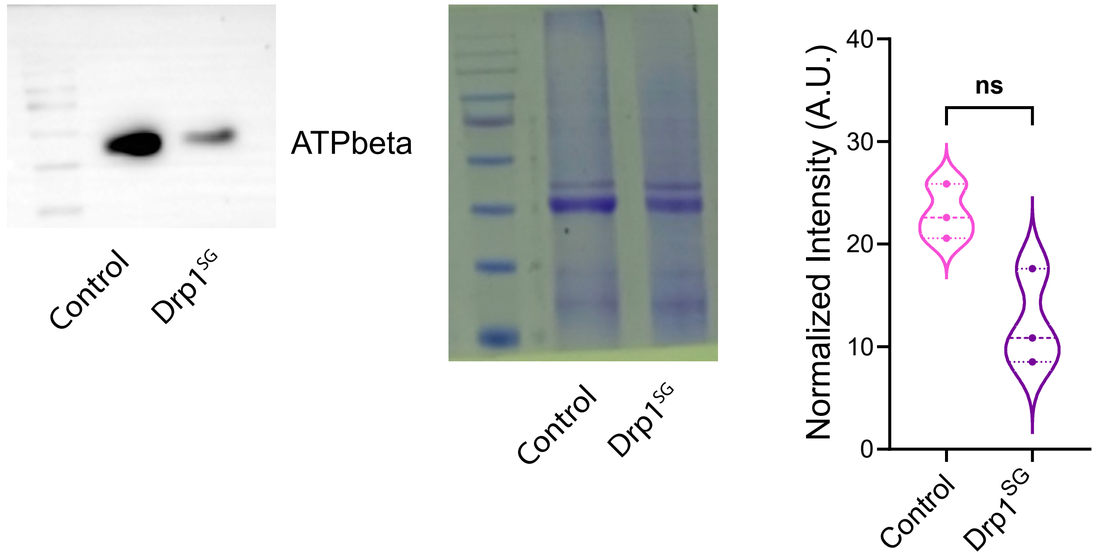
